# A rapid and robust method for single cell chromatin accessibility profiling

**DOI:** 10.1101/309831

**Authors:** Xi Chen, Ricardo J Miragaia, Kedar Nath Natarajan, Sarah A Teichmann

## Abstract

The assay for transposase-accessible chromatin using sequencing (ATAC-seq) is widely used to identify regulatory regions throughout the genome. However, very few studies have been performed at the single cell level (scATAC-seq) due to technical challenges. Here we developed a simple and robust plate-based scATAC-seq method, combining upfront bulk Tn5 tagging with single-nuclei sorting. We demonstrated that our method worked robustly across various systems, including fresh and cryopreserved cells from primary tissues. By profiling over 3,000 splenocytes, we identify distinct immune cell types and reveal cell type-specific regulatory regions and related transcription factors.

---

Due to its simplicity and sensitivity, ATAC-seq^1^ has been widely used to map open chromatin regions across different cell types in bulk. Recent technical developments have allowed chromatin accessibility profiling at the single cell level (scATAC-seq) and revealed distinct regulatory modules across different cell types within heterogeneous samples^2–9^. In these approaches, single cells are first captured by either a microfluidic device^3^ or a liquid deposition system^7^, followed by independent tagmentation of each cell. Alternatively, a combinatorial indexing strategy has been reported to perform the assay without single cell isolation^2,4,9^. However, these approaches require either a specially engineered and expensive device, such as a Fluidigm C1^3^ or Takara ICELL8^7^, or a large quantity of customly modified Tn5 transposase^2,4,5,9^.

Here, we overcome these limitations by performing upfront Tn5 tagging in the bulk cell population, prior to single nuclei isolation. It has been previously demonstrated that Tn5 transposase-mediated tagmentation contains two stages: (1) a tagging stage where the Tn5 transposome binds to DNA, and (2) a fragmentation stage where the Tn5 transposase is released from DNA using heat or denaturing agents, such as sodium dodecyl sulfate (SDS)^10–12^. Since the Tn5 tagging does not fragment DNA, we reasoned that the nuclei would remain intact after incubation with the Tn5 transposome in an ATAC-seq experiment. Based on this idea, we developed a simple, robust and flexible plate-based scATAC-seq protocol, performing a Tn5 tagging reaction^6,13^ on a pool of cells (5,000 – 50,000) followed by sorting individual nuclei into plates containing lysis buffer.

Tween-20 is subsequently added to quench the SDS in the lysis buffer^14^, which otherwise will interfere the downstream reactions. Library indexing and amplification are done by PCR, followed by sample pooling, purification and sequencing. The whole procedure takes place in one single plate, without any intermediate purification or plate transfer steps (Fig. 1 a). With this easy and quick workflow, it only takes a few hours to prepare sequencing-ready libraries, and the method can be implemented by any laboratory using standard equipment.

**Figure 1.**
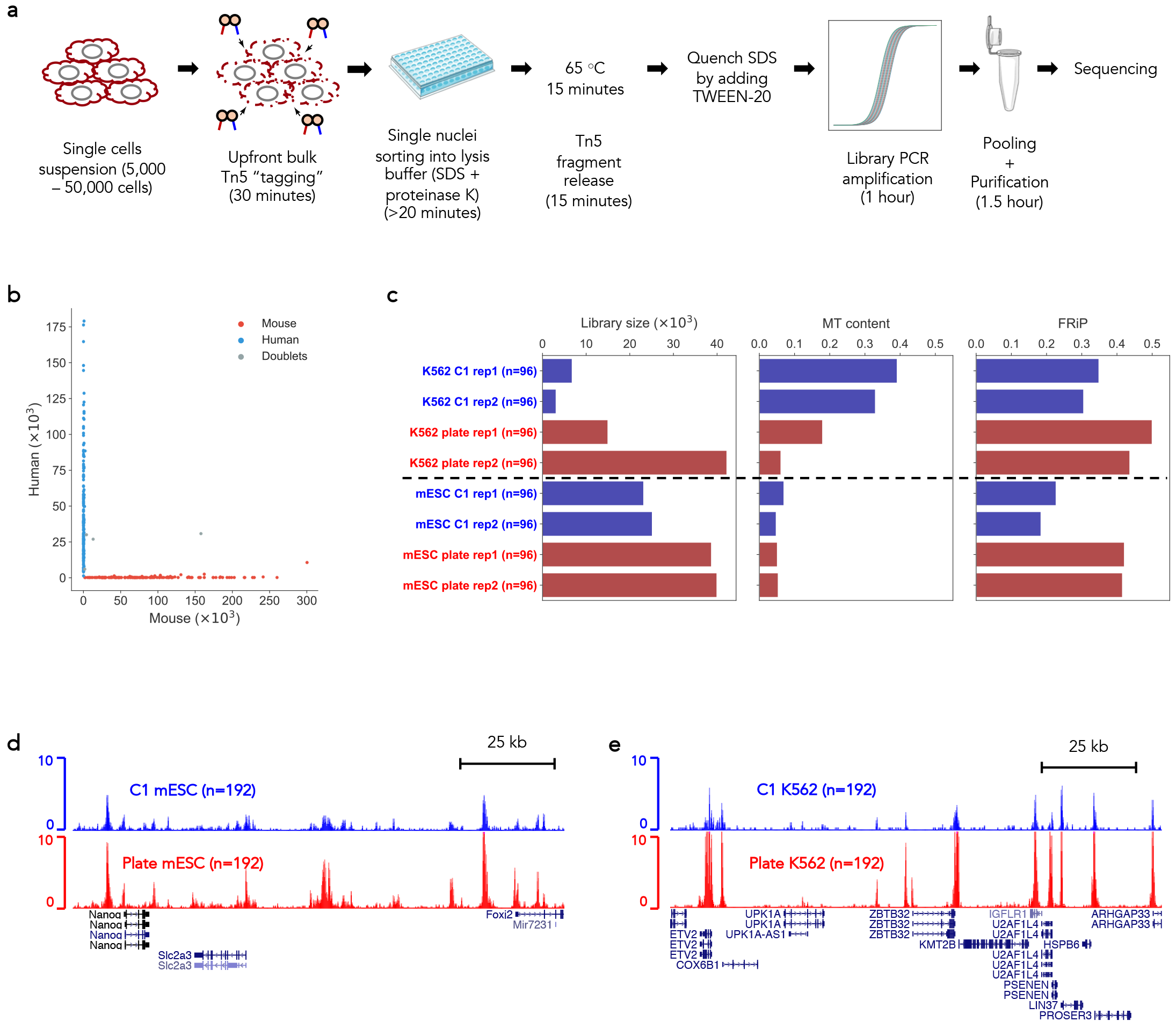
Simple and robust analysis of chromatin status at the single cell level. (a) Schematic view of the workflow of the scATAC-seq method. Tagmentation is performed upfront on bulk cell populations, followed by sorting single-nuclei into 96/384 well plates containing lysis buffer. The lysis buffer contains a low concentration of proteinase K and SDS to denature the Tn5 transposase and fragment the genome. Tween-20 is added to quench SDS^14^. Subsequently, library preparation by indexing PCR is performed, and the number of PCR cycles needed to amplify the library is determined by quantitative PCR (qPCR) (Supplementary Fig. 2b). (b) Species mixing experiments to show the accuracy of FACS. Equal amounts of HEK293T (Human) and NIH3T3 (Mouse) cells were mixed, and scATAC-seq was performed as described in (a). Successful wells with more than 90% of reads uniquely mapped to either human or mouse were categorised as singlets (n=303). Otherwise, they will be categorised as doublets (n=4) (see Online Methods). (c) Comparison of the median library size (estimated by the Picard tool), fraction of mitochondrial DNA (MT content) and fraction of reads in peaks (FRiP) in single cells from either C1 (blue) or plate-based (red) scATAC-seq approach. (d) UCSC genome browser tracks displaying the signal around the *Nanog* gene locus from the aggregate of mESCs obtained from Fluidigm C1 (top) and plate (bottom). (e) The same type of tracks as (d) around the *ZBTB32* gene locus in K562 cells.

We first tested the accuracy of our sorting by performing a species mixing experiment, where equal amounts of HEK293T and NIH3T3 cells were mixed, and scATAC-seq was performed with our method. Using a stringent cut-off (Online Methods), we recovered 307 wells, among which 303 wells contain predominantly either mouse fragments (n = 136) or human fragments (n = 167). Only 4 wells are categorised as doublets (Fig. 1b).

To compare our plate-based method to the existing Fluidigm C1 scATAC-seq approach, we performed side-by-side experiments, where cultured K562 and mouse embryonic stem cells (mESC) were tested by both approaches. We used three metrics to evaluate the quality of the data generated by both methods (Fig. 1 c and Supplementary Fig. 1a). Our plate-based method has higher library complexity (library size estimated by the Picard tool), comparable or lower amount of mitochondrial DNA, and higher signal-to-noise ratio measured by fraction of reads in peaks (FRiP) (Fig. 1 c). In addition, visual inspection of the read pileup from the aggregated single cells suggested both methods were successful, but data generated from our plate-based method exhibited higher signal (Fig. 1d and e).

The main difference between our method and Fluidigm C1 is the Tn5 tagging strategies. The plate-based method performed Tn5 tagging using a population of cells, while it was done in individual microfluidic chambers in the Fluidigm C1. It is possible that the upfront Tn5 tagging is more efficient than tagging in microfluidic chambers.

To evaluate the generality of our method, we tested the plate-based method on cryopreserved cells from four tissues: human and mouse skin fibroblasts (hSF and mSF)^15^ and mouse cardiac progenitor cells (mCPC) at embryonic day E8.5 and E9.5^16^. Cells were revived from liquid nitrogen, and our plate-based method was carried out immediately after revival. The library complexities varied among cell types (Fig. 2 a). We obtained median library sizes ranging from 52,747 (mSF) to 104,608.5 (mCPC_E8.5) unique fragments (Fig. 2a). The amount of mitochondrial DNA also varied across cell types but was low in all samples (<13%). All four tested samples had very high signal-to-noise ratio, with a median FRiP ranging from 0.50 (mSF) to 0.60 (hSF) (Fig. 2 a). The insert size distributions of the aggregated single cells from all four samples exhibited clear nucleosomal banding patterns (Fig. 2 b), which is a feature of high quality ATAC-seq libraries^1^. Finally, visual inspection of aggregate of single cell profiles showed clear open chromatin peaks around expected genes (Fig. 2 c and d). Details of all tested cells/tissues are summarised in Supplementary Table 1.

**Figure 2.**
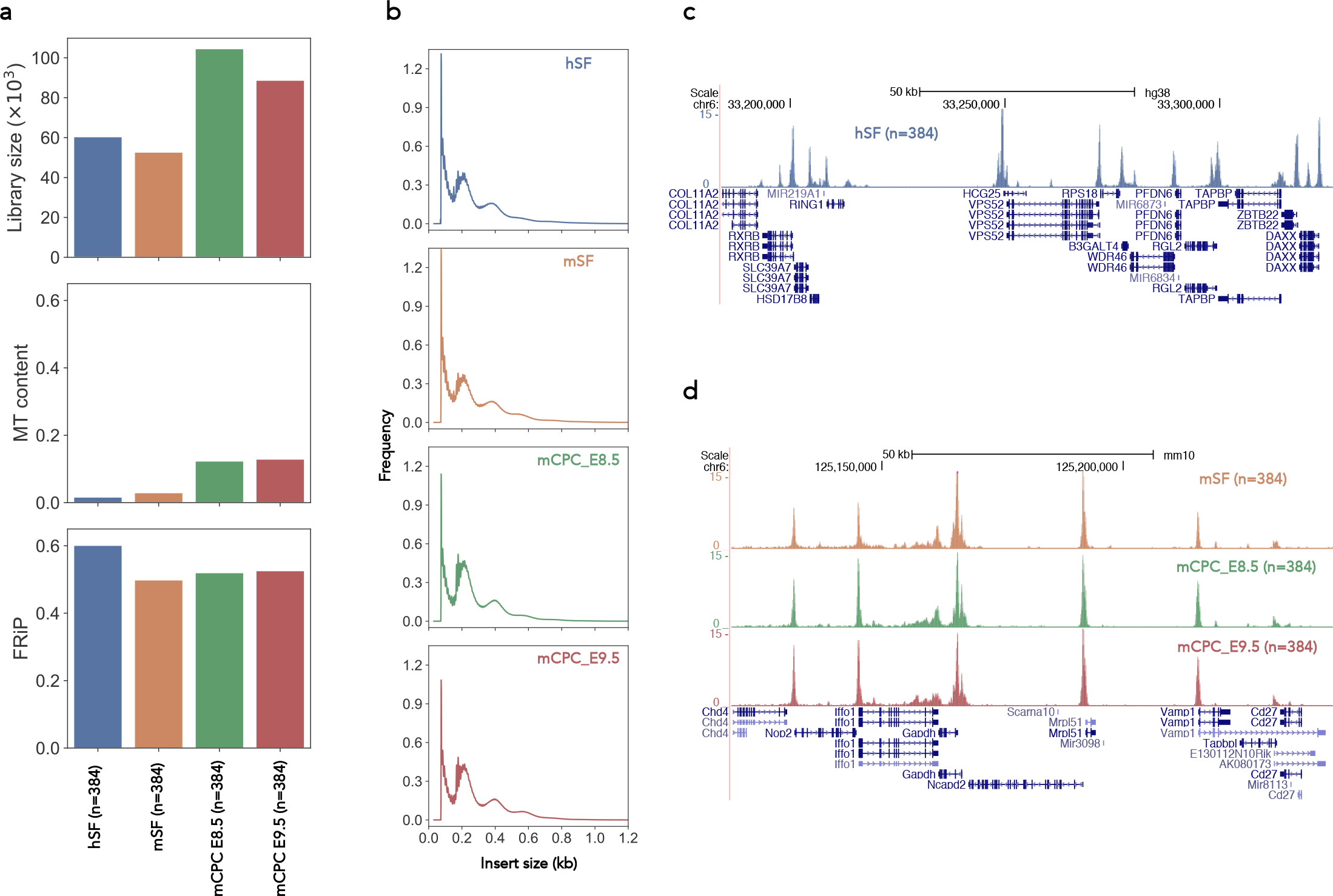
Plate-based scATAC-seq worked robustly on cryopreserved cells from primary tissues. (a) Comparison of the median library size (estimated by the Picard tool), fraction of mitochondrial DNA (MT content) and fraction of reads in peaks (FRiP) in cryopreserved single cells from four different tissues: human skin fibroblasts (hSF), mouse skin fibroblasts (mSF), mouse cardiac progenitor cells (mCPC) at embryonic day E8.5 and E9.5. (b) Insert size frequencies from the aggregated data of the cells from the four tissues. (c and d) UCSC genome browser tracks displaying the signal around the *RPS18* gene locus from the aggregate of hSFs (c) and around the *Gapdh* gene locus from the aggregate of mSFs, mCPC_E8.5 and mECP_E9.5 (d).

After this validation of the technical robustness of our plate-based method, we further tested it by generating the chromatin accessibility profiles of 3,648 splenocytes (after red blood cell removal) from two C57BL/6Jax mice. In total, we performed two 96-well plates and nine 384-well plates. By setting a stringent quality control threshold (>10,000 reads and >90% mapping rate), 3,385 cells passed the technical cut-off (>90% successful rate) (Supplementary Fig. 3b). The aggregated scATAC-seq profiles exhibited good coverage and signal and resembled the bulk data generated from 10,000 cells by the Immunological Genome Project (ImmGen)^17^ (Fig. 3a). The library fragment size distribution before and after sequencing both displayed clear nucleosome banding patterns (Fig. 3 b and Supplementary Fig. 2 a). In addition, sequencing reads showed strong enrichment around transcriptional start sites (TSS) (Fig. 3c), further demonstrating the quality of the data was high.

**Figure 3.**
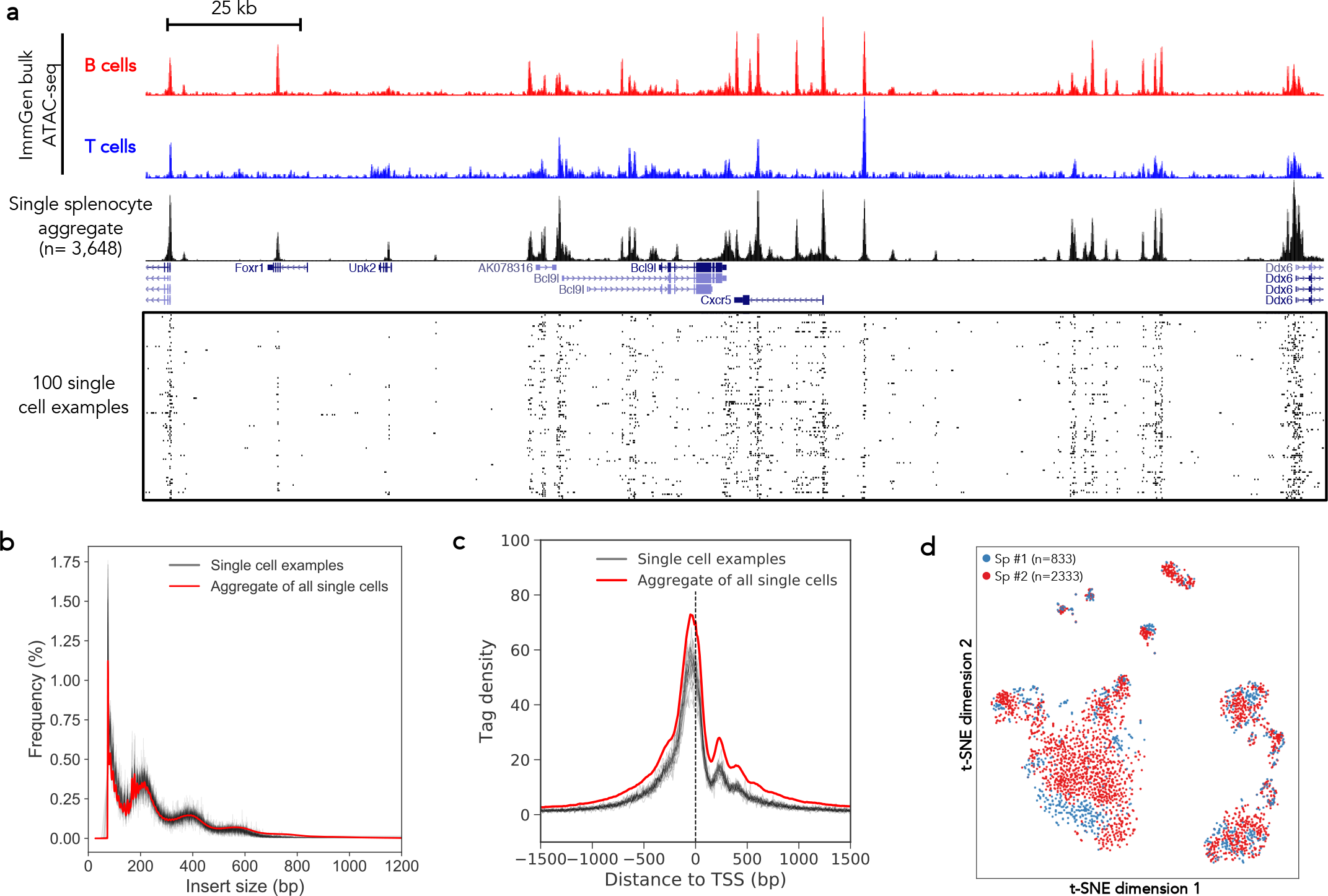
Plate-based scATAC-seq applied to over 3000 mouse splenocytes. (a) UCSC genome browser tracks displaying the signal around the *Cxcr5* gene locus from the aggregate of all single cells in this study. Bulk ATAC-seq profiles from the ImmGen consortium are also shown. Randomly selected 100 single cell profiles are show below the aggregated profile. (b and c) Insert size frequencies (b) and sequencing read distributions across transcriptional start sites (c) of libraries from the aggregated data (the red line) and individual single cells (grey lines, 24 examples are shown). (d) A two-dimensional projection of the scATAC-seq data using t-SNE. Colours represent two different batches, showing excellent agreement between batches. Sp: spleen.

Importantly, for the majority of the cells, less than 10% (median 2.1%) of the reads were mapped to the mitochondrial genome (Supplementary Fig. 3a). Overall, we obtained a median of 643,734 reads per cell, while negative controls (empty wells) generated only ~ 100-1,000 reads (Supplementary Fig. 3b). In most cells, more than 98% of the reads were mapped to the mouse genome (Supplementary Fig. 3b), indicating low level of contamination. The median of estimated library sizes is 31,808.5 (Supplementary Fig. 3c). At the sequencing depth of this experiment, the duplication rate of each single cell library is ~ 95% (Supplementary Fig. 3 d), indicating that the libraries were sequenced to near saturation. Downsampling the raw reads (from the fastq files) and repeating the analysis suggest that at 20 – 30% of our current sequencing depth, the majority of the fragments would have already been captured (Supplementary Fig. 4a and b). Therefore, in a typical scATAC-seq experiment, ~ 120,000 reads per cell are sufficient to capture most of the unique fragments, with higher sequencing depth still increasing the number of detected unique fragments (Supplementary Fig. 3e).

Next, we examined the data to analyse signatures of different cell types in the mouse spleen. Reads from all cells were merged, and a total of 78,048 open chromatin regions were identified by peak calling with q values less than 0.01^18^ (Online Methods). We binarised peaks as “open” or “closed” (Online Methods) and applied a Latent Semantic Indexing (LSI) analysis to the cell-peak matrix for dimensionality reduction^2^ (Online Methods). Consistent with previous findings^2^, the first dimension is primarily influenced by sequencing depth (Supplementary Fig. 3f). Therefore, we only focused on the second dimension and upwards and visualised the data by t-distributed stochastic neighbour embedding (t-SNE)^19^. We did not observe batch effects from the two profiled spleens, and several distinct populations of cells were clearly identified in the t-SNE plot (Fig. 3 d). Read counts in peaks near key marker genes (*e.g. Bcl11a* and *Bcl11b*)suggested that the major populations are B and T lymphocytes, as expected in this tissue (Fig. 4a). In addition, we found a small number of antigen-presenting cell populations (Supplementary Fig. 5), consistent with previous analyses of mouse spleen cell composition^20^.

**Figure 4.**
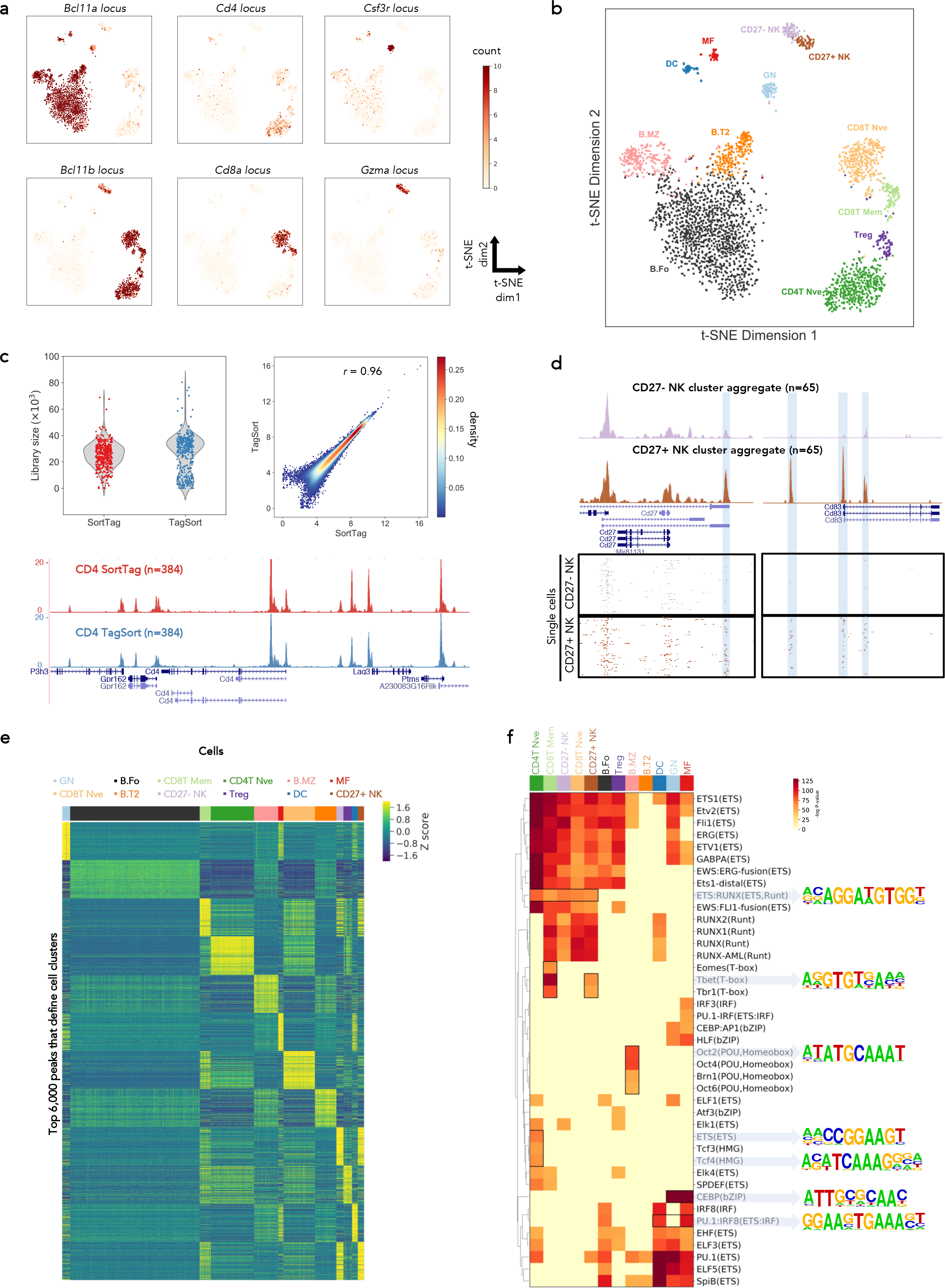
Identification of different cell types and cell-type specific open chromatin regions and transcription factor motifs. (a) The same t-SNE plot as in Figure 3d, coloured by the number of counts in the peaks near indicated gene locus. (b) The same t-SNE plot as in Figure 3d coloured by spectral clustering and cell type annotation. (c) Comparisons of spleen CD4 T cells scATAC-seq obtained by two strategies. TagSort: cells were stained with anti-CD4-PE, tagged with Tn5 and CD4-PE positive cells were sorted for scATAC-seq; SortTag: CD4 T cells were purified first and scATAC-seq was performed on the purified cells. Top: comparison of library size and binding signal correlation (pearson r=0.96) around called peaks; bottom: UCSC genome browser tracks of the indicated single cell aggregates around the *Cd4* gene locus. (d) UCSC genome browser tracks around *Cd27* and *Cd83* gene loci, displaying the aggregate (top panel) and single cell (bottom panel) signals of the two NK clusters. ATAC-seq peaks specific to the CD27+ NK cells are highlighted. For visual comparison reason, we randomly choose 65 out of 75 CD27-NK cells. (e) Z-score of normalised read counts in the top 500 peaks that distinguish each cell cluster based on the logistic regression classifier, across each peak (row) in each cell (column). Top 500 marker peaks were picked per cell cluster, so there are 500 × 12 = 6,000 peaks in the heatmap. Cells are ordered by cluster labels. (f) Heatmap representation of transcription factor motif (rows) enrichments (binomial test p-values) in the top 500 marker peaks in different cell clusters (columns). Some key motifs are enclosed by black rectangles and motif logos are shown to the right. Motif names are taken from the HOMER software suite.

To systematically interrogate various cell populations captured in our experiments, we applied a spectral clustering technique^21^ which revealed 12 different cell clusters (Fig. 4b). Reads from cells within the same cluster were merged together to form ‘pseudo-bulk’ samples and compared to the bulk ATAC-seq data sets generated by ImmGen (Supplementary Fig. 6 and 7). Cell clusters were assigned to the most similar ImmGen cell type (Fig. 4b and Supplementary Fig. 7). In this way, we identified most clusters as different subtypes of B, T and Natural Killer (NK) cells, as well as a small population of granulocytes (GN), dendritic cells (DC) and macrophages (MF) (Fig. 4b and Supplementary Table 2). An aggregate of all single cells within the same predicted cell type agrees well with the ImmGen bulk ATAC-seq profiles (Supplementary Fig. 8). Remarkably, the aggregate of as few as 55 cells (e.g. the predicted MF cell cluster) already exhibited typical bulk ATAC-seq profiles (Supplementary Fig. 8). This finding opens the door for a novel ATAC-seq experimental design, where Tn5 tagging can be performed upfront on large populations of cells (e.g. 5,000 – 50,000 cells). Subsequently, cells of interest (for example, marked by surface protein antibodies or fluorescent RNA/DNA probes) can be isolated by FACS, and libraries generated for subsets of cells only. This will be a simple and fast way of obtaining scATAC-seq profiles for rare cell populations.

To test the feasibility of this idea, we stained mouse splenocytes with an anti-CD4antibody conjugated with PE and performed tagmentation afterwards. The PE signal remained after tagmentation (Supplementary Fig. 9), allowing us to specifically sort out CD4 positive T cells from the rest of the splenocytes for analysis (we named these “TagSort” libraries). As a control, we first purified CD4 T cells using an antibody-based depletion method (Online Method), and subsequently performed scATAC-seq on the purified CD4 T cells (we named these “SortTag” libraries). The data of CD4 T cells generated from these two strategies agree very well (Fig. 4c). The library complexity is comparable with median library sizes of 30,953 and 25,830 respectively (Fig. 4c, top left panel). The binding signals around open chromatin peaks are highly correlated (Pearson r=0.96) (Fig. 4c, top right panel). Visual inspection of read pileup profiles around the *Cd4* gene locus from single cell aggregates suggested the data are of good quality (Fig. 4c, bottom panel).

This experiment serves as a proof-of-principle test where staining of a surface marker can be done before Tn5 tagging, and a specific population can be sorted by FACS afterwards for scATAC-seq analysis. It should be noted that we have only tested CD4 - an abundant marker in a subpopulation of splenocytes. Other surface markers in different tissues would need to be investigated individually. In addition, the ability to investigate rare cell populations using this approach is limited by the frequency of the rare cell types and the amount of cells that can be tagged upfront.

The spectral clustering was able to distinguish different cell subtypes, such as naïve and memory CD8 T cells, naïve and regulatory CD4 T cells and CD27+ and CD27− NK cells (Fig. 4b). Previous studies have identified many enhancers that are only accessible in certain cell subtypes, and these are robustly identified in our data. Examples are the *Ilr2b* and *Cd44* loci in memory CD8 T cells^22^ and *Ikzf2* and *Foxp3* in regulatory T cells^23^ (Supplementary Fig. 10a and b). Interestingly, our clustering approach successfully identified two subtle subtypes of NK cells (CD27− and CD27 + NK cells), as determined by their open chromatin profiles (Fig. 4b and d). It has been shown that, upon activation, NK cells can express CD83^24^, a well-known marker for mature dendritic cells^25^. In mouse spleen, *Cd83* expression was barely detectable in the two NK subpopulations profiled by the ImmGen consortium (Supplementary Fig. 10 c). However, in our data, the *Cd83* locus exhibited different open chromatin states in the two NK clusters (Fig. 4 d). Multiple ATAC-seq peaks were observed around the *Cd83* locus in the CD27+ NK cell cluster but not in the CD27-NK cluster (Fig. 4d). This suggests that *Cd83* is in a transcriptionally permissive state in the Cd27+ NK cells, and the CD27+ NK cells have a greater potential for rapidly producing CD83 upon activation. This may partly explain the functional differences between CD27 + and CD27− NK cell states^26^.

Finally, we investigated whether we could identify the regulatory regions that define each cell cluster. To this end, we trained a logistic regression classifier using the spectral clustering labels and the binarised scATAC-seq count data (Online Methods). From the classifier, we extracted the top 500 open chromatin peaks (marker peaks) that can distinguish each cell cluster from the others (Fig. 4 e and Online Methods). By looking at genes in the vicinity of the top 50 marker peaks, we recapitulated known markers, such as *Cd4* for the helper T cell cluster (cluster 3), *Cd8a* and *Cd8b1* for the cytotoxic T cell cluster (cluster 6) and *Cd9*for marginal zone B cell cluster (cluster 4) (Supplementary Figure 11 and Supplementary Table 3). These results are consistent with our correlation-based cell cluster annotation (Fig. 4b).

While the peaks at TSS are useful for cell type annotation, the majority of the cluster-specific marker peaks are in intronic and distal intergenic regions, in line with the global peak distribution (Supplementary Fig. 12). To identify transcription factors that are important for the establishment of these marker peaks, we investigated them in more detail by motif enrichment analysis using HOMER^27^. The full results of these motif enrichment analyses are included in Supplementary Table 4. As expected, different ETS motifs and ETS-IRF composite motifs were significantly enriched in marker peaks of many clusters (Fig. 4f), consistent with the notion that ETS and IRF transcription factors are important for regulating immune activities^28^. Furthermore, we found motifs that were specifically enriched in certain cell clusters (Fig. 4f). Our motif discovery is consistent with previous findings, such as the importance of T-box (*e.g.* Tbx21) motifs in NK^29^ and CD8T memory cells^30^ and POU domain (e.g. Pou2f2) motifs in marginal zone B cell^31^. This suggests that our scATAC-seq data is able to identify known gene regulation principles in different cell types within a tissue.

In recent years, other methods, such as DNase-seq^32^, MNase-seq^33^ and NOMe-seq^34,35^, have investigated chromatin status at the single cell level. However, due to its simplicity and reliability, ATAC-seq currently remains the most popular technique for chromatin profiling. Several recent studies have demonstrated the power of using scATAC-seq for investigating regulatory principles, *e.g.* brain development^4,9^, Mouse sci-ATAC-seq Atlas^36^ and pseudotime inference^37^. The combined multi-omics approaches also began to emerge, such as sci-CAR-seq^38^, scCAT-seq^39^ and piATAC-seq^8^. Our study added on top of those methods to provide a simple and easy-to-implement scATAC-seq approach that can successfully detect different cell populations, including subtle and rare cell subtypes, from a complex tissue. More importantly, it is able to reveal key gene regulatory features, such as cell-type specific open chromatin regions and transcription factor motifs, in an unbiased manner. Future studies can utilise this method to unveil the regulatory characteristics of novel and rare cell populations and the mechanisms behind their transcriptional regulation.

## Author contributions

X.C, K.N.N and S.A.T conceived the project. X.C designed the protocol. X.C, R.J.M and K.N.N performed the experiments. X.C carried out the computational analysis. S.A.T supervised the entire project. All authors contributed to the writing.

## Acknowledgement

We thank Jong-Eun Park, Johan Henriksson, Tzachi Hagai, Tomas Gomez, Kerstin Meyer, Roser Vento, Lira Mamanova and all others from the Teichmann group for the inspiring discussion of the method, the critical reading of the manuscript and the computational help. We also thank Natalia Kunowska, Qianxin Wu and Andrew Knights for the helpful discussion related to the Tn5 transposase. We thank Bee Ling Ng, Chris Hall, Jennie Graham and Sam Thompson for the excellent support of FACS. We thank the DNA pipeline from the Wellcome Sanger Institute for the Illumina sequencing support. We thank Guangshuai Jia and Jens Preussner from Thomas Braun’s lab for sharing the mouse cardiac progenitor cells. We thank Aik Ooi for the initial help with the experimental setup.

## Data and materials availability

The sequencing data has been deposited at ArrayExpress (accession E-MTAB-6714). The code used for the analysis is available on the Github repository https://github.com/dbrg77/plate_scATAC-seq.

The UCSC genome browser tracks containing both the ImmGen bulk ATAC-seq and scATAC-seq from this study can be viewed via this link: http://genome-euro.ucsc.edu/cgi-bin/hgTracks?hgS_doOtherUser=submit&hgS_otherUserName=dbrg77&hgS_otherUserSessionName=mSpleen_scATAC_cluster.

## Methods

### Cell isolation

For splenocytes, the spleen from a C57BL/6Jax mouse was mashed by a 2-ml syringe plunger through a 70 μm cell strainer (Fisher Scientific 10788201) into 30 ml 1X DPBS (ThermoFisher 14190169) supplied with 2 mM EDTA and 0.5% (w/v) BSA (Sigma A9418). Cells were centrifuged down, supernatant was removed, and the cell pellet was briefly vortexed. 5 ml 1X RBC lysis buffer (ThermoFisher 00-4300-54) was used to resuspend the cell pellet, and the cell suspension was vortexed again, and left on bench for 5 minutes to lyse red blood cells. Then 45 ml 1X DPBS was added, and cells were centrifuged down. 30 ml 1X DPBS were used to resuspend the cell pellet. The cell suspension was passed through a Miltenyi 30 μm Pre-Separation Filter (Miltenyi 130-041-407), and the cell number was determined using C-chip counting chamber (VWR DHC-N01). All centrifugations were done at 500 g, 4 °C, 5 minutes. For human and mouse skin fibroblasts, cells were extracted as previously described^15^. For mouse cardiac progenitor cells, cells were extracted as previously described^16^. Cells were cryopreserved in 90% FBS and 10% DMSO and stored in liquid nitrogen until experiments.

### Plate-based single-cell ATAC-seq (scATAC-seq)

A detailed step-by-step protocol can be found in the Supplementary Protocol. Briefly, 50,000 cells were centrifuged down at 500 g, 4 °C, 5 minutes. Cell pellets were resuspended in 50 μl tagmentation mix (33 mM Tris-acetate, pH 7.8, 66 mM potassium acetate, 10 mM magnesium acetate, 16% dimethylformamide (DMF), 0.01% digitonin and 5 μl of Tn5 from the Nextera kit from Illumina, Cat. No. FC-121-1030). The tagmentation reaction was done on a thermomixer (Eppendorf 5384000039) at 800 rpm, 37 °C, 30 minutes. The reaction was then stopped by adding equal volume (50 μl) of tagmentation stop buffer (10 mM Tris-HCl, pH 8.0, 20 mM EDTA, pH 8.0) and left on ice for 10 minutes. 200 μl 1X DPBS with 0.5% BSA was added and the nuclei suspension was transferred to a FACS tube. DAPI (ThermoFisher 62248) was added at a final concentration of 1 μg/μl to stain the nuclei.

### Species mixing experiments

25,000 HEK293T (Human) and 25,000 NIH3T3 (Mouse) cells were mixed together, and scATAC-seq was performed as described in the Supplementary Protocol. The obtained sequencing reads were mapped to a concatenated genome of mouse and human by hisat2^40^. One 384-well plate was performed. We first set a technical cutoff where a successful well must contain more than 10,000 total reads and more than 90% of reads are mapped to the concatenated genome. 307 wells were marked as successful. Among the successful wells, we calculated the ratio of reads that mapped to the human genome and the mouse genome. If the ratio is larger than 10, the well is categorised as containing human single cells; if the ratio is less than 0.1, the well is categorised as containing mouse single cells; otherwise, the well is categorised as containing human-mouse doublets.

### Plate scATAC-seq on CD4+ T cells (TagSort vs SortTag)

For the “TagSort” strategy, 50,000 splenocytes were stained with anti-Mouse CD4 PE (eBioscience cat no. 12-0043-82) at room temperature for 30 minutes according to the manufacturer’s instructions. The stained cells were washed with ice-cold 1X PBS twice and pelleted down at 500 g, 4 °C, 5 minutes. Experimente were carried out following the procedures described in the Supplementary Protocol. DAPI and PE double positive cells were sorted into a 384-well plate for library construction. For the “SortTag” strategy, CD4+ T cells were purified first from mouse splenocytes using the Naive CD4 T Cell Isolation Kit, Mouse (Miltenyi, cat. no. 130-104-453) following the manufacturer’s instruction without the anti-CD44 depletion step. The purified CD4 T cells were processed according to the procedures described in the Supplementary Protocol.

### scATAC-seq using Fluidigm C1

Experiments were performed as previously described^3^ using the medium-sized (1862×) Open App chip. We followed the manufacturer’s instructions described in the “ATAC Seq No Stain (Rev C)” from the Fluidigm ScriptHub (https://www.fluidigm.com/c1openapp/scripthub), except that we replace the detergent NP-40 in the original protocol with digitonin so that the final concentration of digitonin in the reaction chamber is 0.005%. After collecting the pre-amplified material from the Fluidigm chip, the libraries were indexed by library PCR for 14cycles as previously described^3^.

### Costs involved in plate-based and Fluidigm C1 scATAC-seq

For our plate-based scATAC-seq method, most reagents and buffers are available in a standard molecular biology lab. Exceptions are the Tn5 transposase, which can be purchased from Illumina (Cat No. FC-121-1030), and the PCR master mix, which can be purchased from various vendors (we used the 2X NEBNext^®^ High-Fidelity 2X PCR Master Mix from NEB). Since the Tn5 tagging reaction was performed upfront at the bulk level, the Tn5 cost per cell depends on how many cells are sorted during the sorting. Based on our experience, when 50,000 cells are used at the beginning, two to eight 384-well plates can be sorted. Therefore, the cost of Tn5 is negligible. The major cost per unit for the plate-based scATAC-seq is the PCR master mix used during library amplification. Currently, 10 μl of PCR master mix are needed per cell in a 20 μl library amplification reaction, but we have been successfully and consistently generated libraries from half of the volume described in the protocol. For scATAC-seq using the Fluidigm C1, all the aforementioned reagents are needed, and a microfluidic chip is required per 96 cells.

### Hands-on time for plate-based versus C1 scATAC-seq approaches

For our plate-based scATAC-seq method, the most time-consuming part is the lysis plate preparation (mixing lysis buffer and indexing primers). For maximum efficiency, this can be done upfront in bulk, and the lysis plate is stable in −80 °C for a long time. Another time/labour-consuming step is the pooling of single cell libraries after PCR using a multi-channel pipette. We provide online advice to perform the whole procedure in minutes. This information is included in the accompanying GitHub page: https://github.com/dbrg77/plate_scATAC-seq. For scATAC-seq using the Fluidigm C1, an extra ~4 hours of C1 runtime are needed.

### qPCR for library amplification

After assembly of the 20 μl PCR reaction (see Supplementary Protocol), a pre-amplification step was performed on a PCR machine (Alpha Cycler 4, PCRmax) with 72 °C 5 minutes, 98 °C 5 minutes, 8 cycles of [98 °C 10 seconds, 63 °C 30 seconds, 72 °C 20 seconds]. Of the product, 19 μl of pre-amplified library were transferred to a 96 well qPCR plate, 1 μl 20X EvaGreen (Biotium #31000) was added, and qPCR was performed on an ABI StepOnePlus system with the following cycle conditions: 98 °C 1 minutes, 20 cycles of [98 °C 10 seconds, 63 °C 30 seconds, 72 °C 20 seconds]. Data was acquired at 72 °C. We qualitatively chose the cycle number to where the fluorescence signals just about to start going up (Supplementary Fig. 1b). In this study, a total of 18 cycles were used to amplify the libraries.

### Sequencing data processing

All sequencing data were processed using a pipeline written in snakemake^41^. The software/packages and the exact flags used in this study can be found in the ‘Snakefile’ provided in the GitHub repository https://github.com/dbrg77/plate_scATAC-seq. Briefly, reads were trimmed with cutadapt^42^ to remove the Nextera sequence at the 3′ end of short inserts. The trimmed reads were mapped to the reference mouse genome (UCSC mm10) using hisat2^40^. Reads with mapping quality less than 30 were removed by samtools^43^ (−q 30 flag) and deduplicated using the MarkDuplicates function of the Picard tool (http://broadinstitute.github.io/picard). All reads from single cells were merged together using samtools, and the merged BAM file was deduplicated again. Peak calling was performed on the merged and deduplicated BAM file by MACS2^18^. For bulk ATAC-seq and single cell aggregate coverage visualisation, bedGraph files generated from MACS2 callpeak were converted to bigWig files and visualised via UCSC genome browser. For individual single cell ATAC-seq visualisation, aligned reads from individual cells were converted to bigBed files. A count matrix over the union of peaks was generated by counting the number of reads from individual cells that overlap the union peaks using coverageBed from the bedTools suite^44^.

### Public ATAC-seq data processing

FASTQ files were all downloaded from the European Nucleotide Archive (ENA). The ImmGen bulk ATAC-seq data (study accession PRJNA392905) and the scATAC-seq data using Fluidigm C1 (study accessions PRJNA274006 and PRJNA299657) were processed in the same way as described in this study. The ‘Snakefile’ used to process the data can be found at the the same GitHub repository.

### Bioinformatics analysis

Codes used to carry out all the analyses were provided as Jupyter Notebook files, which can be found in the same GitHub repository. Briefly, downsampling was performed by randomly selecting a fraction of reads from the original FASTQ files using seqtk (https://github.com/lh3/seqtk), and the same pipeline was run on the sub-sampled FASTQ files. For binarising the scATAC-seq data, peak calling was performed on reads merged from all cells, and we labelled the peak ‘1’ (open) if there was at least one read overlapping the peak, and ‘0’ (closed) otherwise. Latent semantic indexing analysis was performed by first normalising the binarized count matrix by term frequency inverse document frequency (TF-IDF) and then performing a Singular-Value Decomposition (SVD) on the normalised count matrix. Only the 2nd – 50th dimensions after the SVD were passed to t-SNE for visualisation. To compare with ImmGen bulk ATAC-seq data, a reference peak set was created by taking the union of peaks from the peak calling results of aggregated scATAC-seq (this study) and different samples of ImmGen bulk ATAC-seq using mergeBed from the bedTools suite^44^. All comparisons were done based on this reference peak set. The annotatePeaks.pl from HOMER^27^ was used to assign genes to peaks. Latent semantic indexing, spectral clustering and logistic regression were carried out using Scikit-learn^45^.

---

**Supplementary Figure 1.**
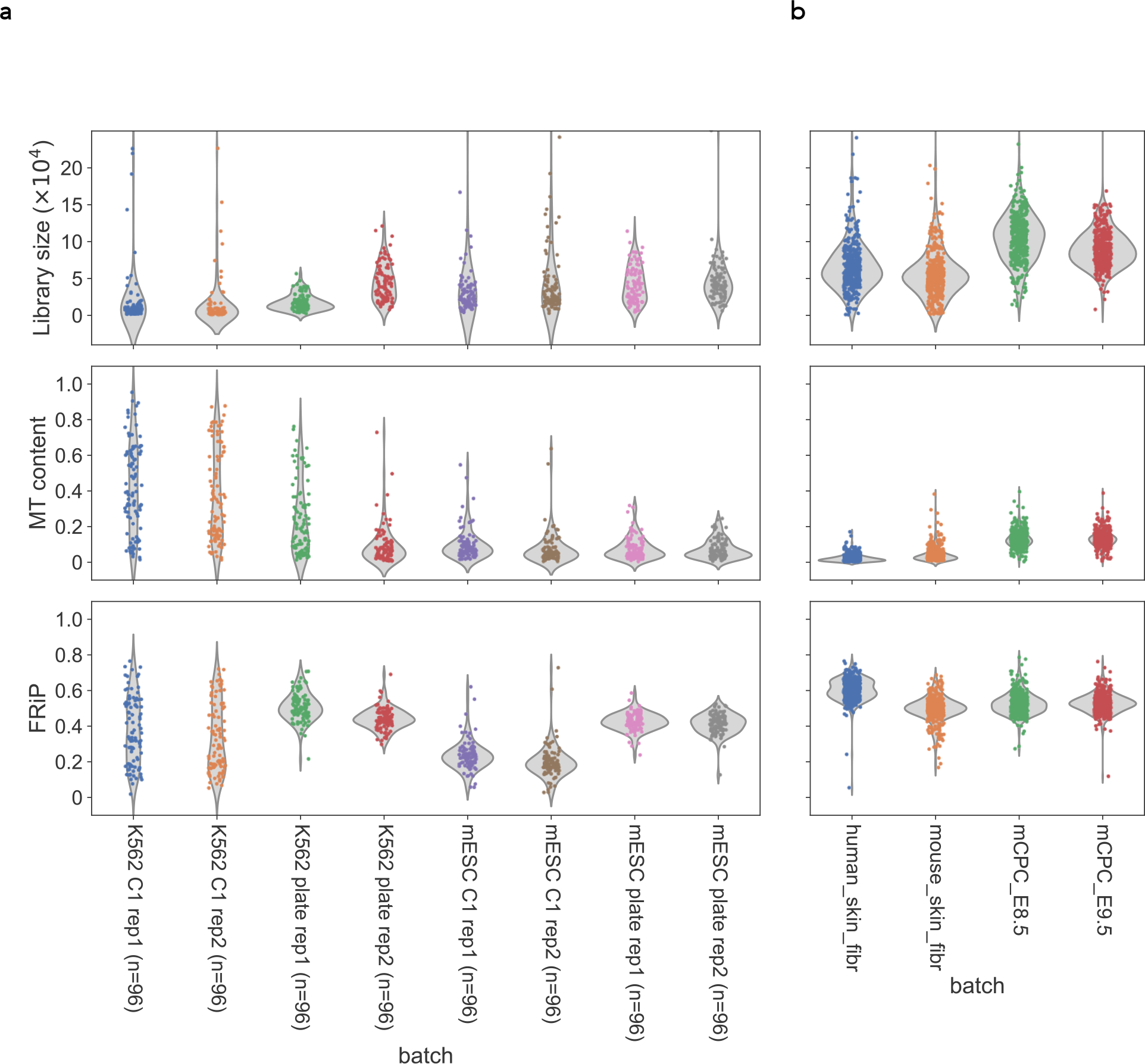
(a) Violin plots showing the comparisons of distributions of the median library size (estimated by the Picard tool), fraction of mitochondrial DNA (MT content) and fraction of reads in peaks (FRiP) in single cells from either plate or C1 scATAC-seq approach. (b) the same metrics as in (a) showing data obtained from cryopreserved cells of four different primary tissues.

**Supplementary Figure 2.**
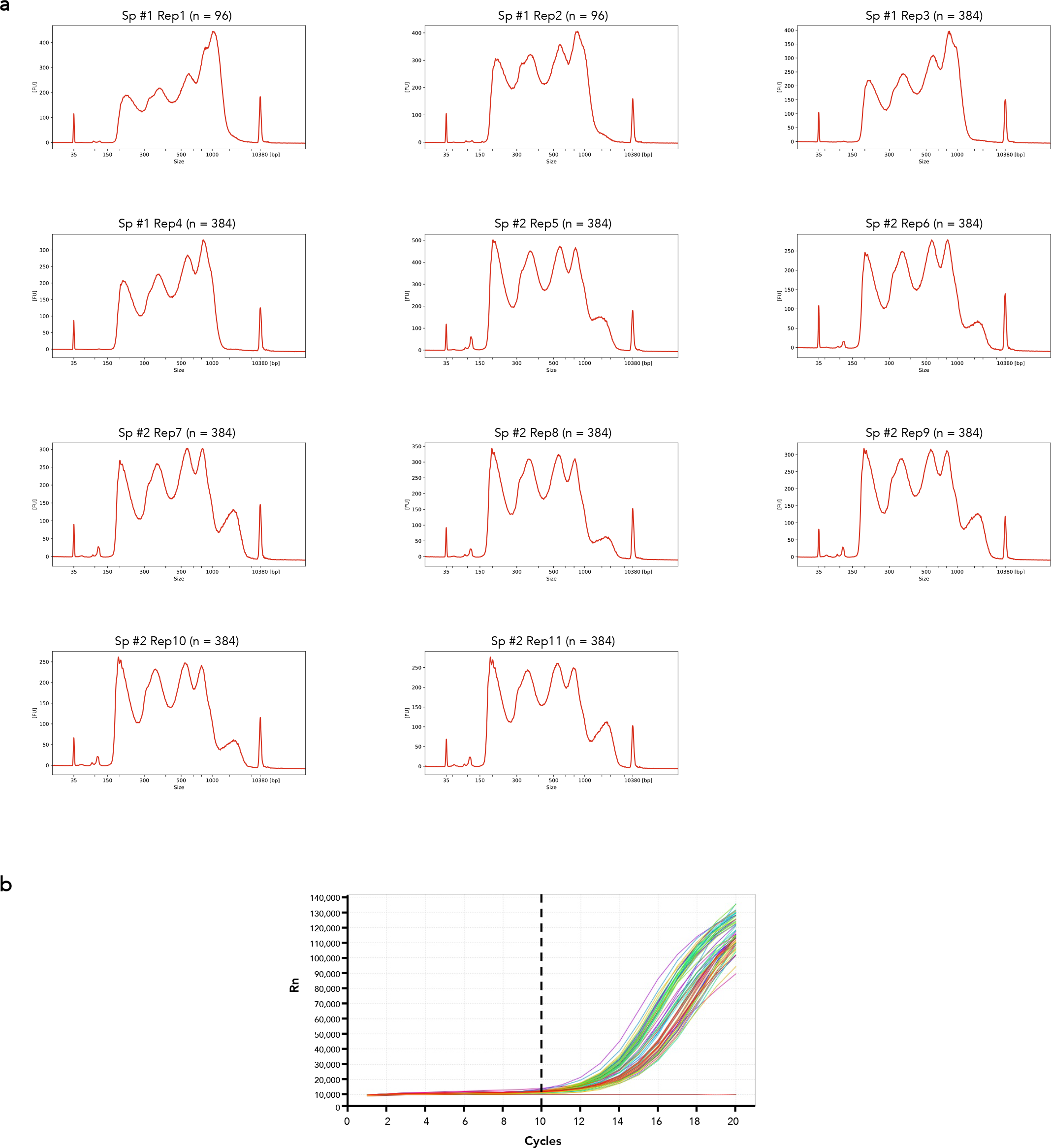
(a) Bioanalyzer results of pools of 11 different plates (two spleens) of scATAC-seq in this study. (b) qPCR amplification plot of 64 different single cell libraries. The qPCR was performed after 8 cycles of pre-amplification. Dotted line indicates the number of cycles used for final amplification. A total of 8 + 10 = 18 cycles were performed in this study.

**Supplementary Figure 3.**
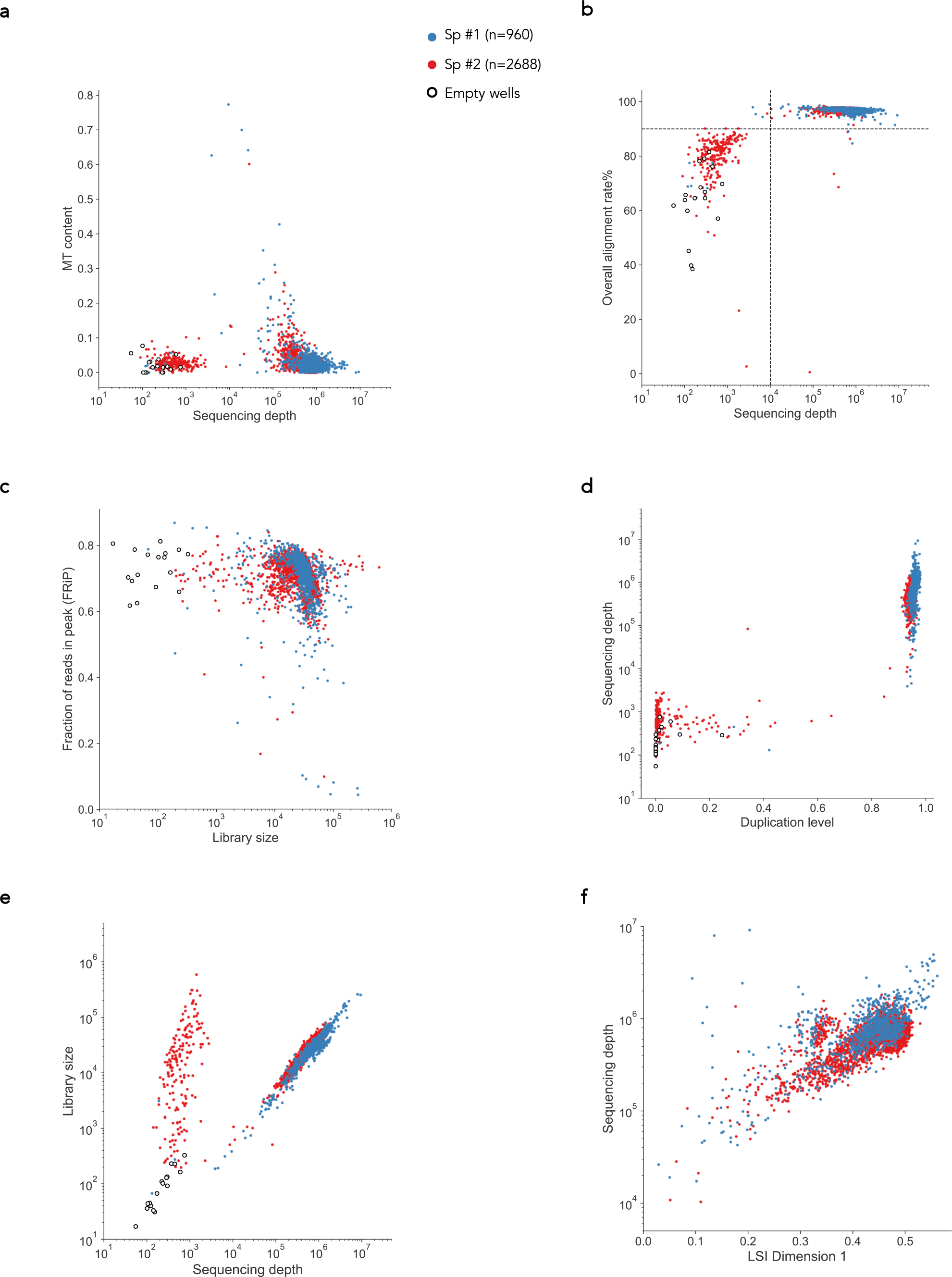
Scatter plots of different quality control metrics. Single cells from different batches are indicated by different colours, and empty well controls are also indicated. We removed cells that have less than 10,000 reads or less than 90% mapping rate, as indicated by dotted lines in (b). Sp: spleen; MT content: fraction of mitochondrial reads; LSI: latent semantic indexing; Library size is estimated by the Picard tool.

**Supplementary Figure 4.**
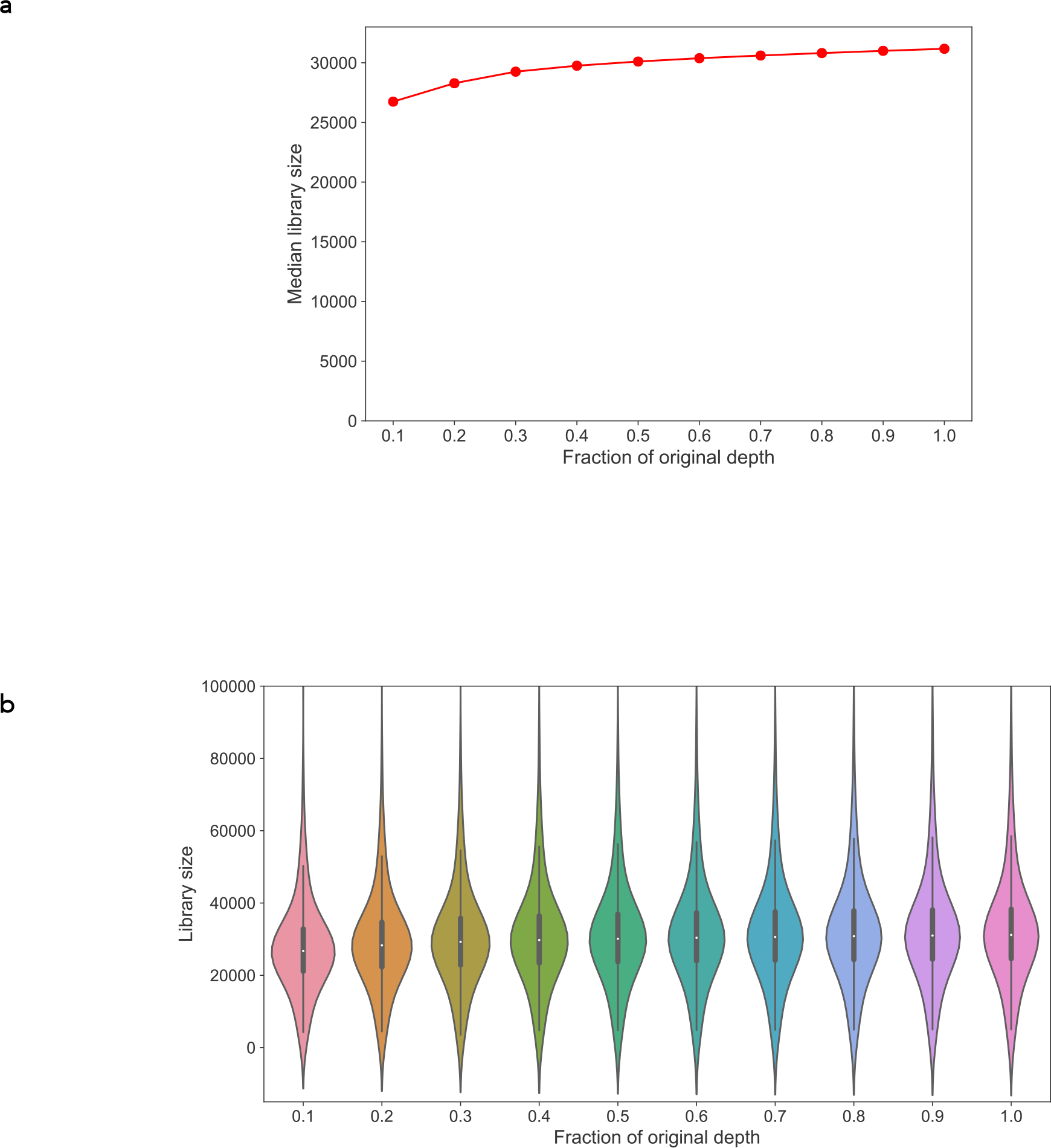
(a) The median library size after downsampling (at the fastq stage) to different fractions relative to the full data sets. (b) Violin Plot of the library size at the different level of downsampling.

**Supplementary Figure 5.**
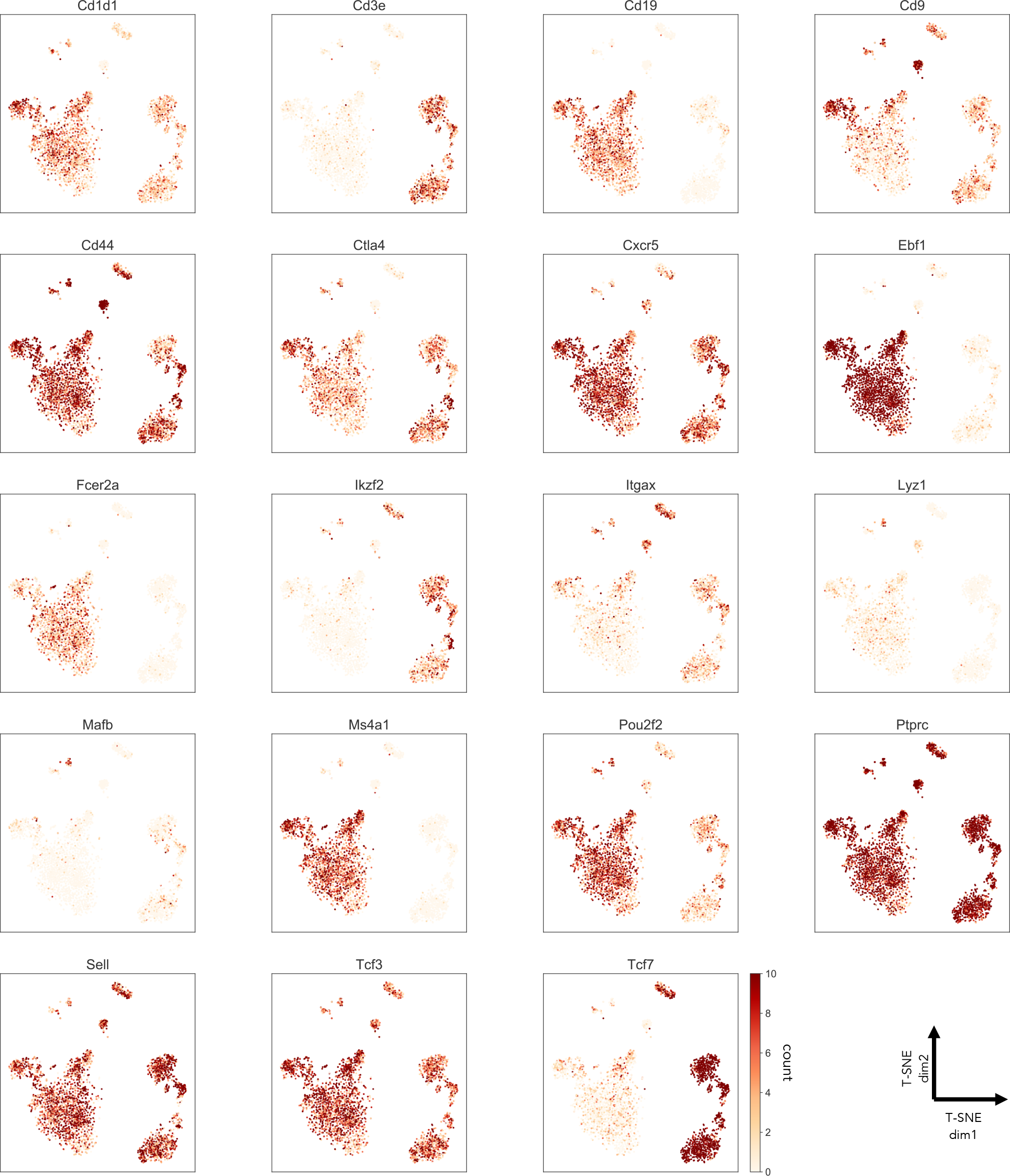
Number of counts from all peaks that assigned to the indicated genes by HOMER.

**Supplementary Figure 6.**
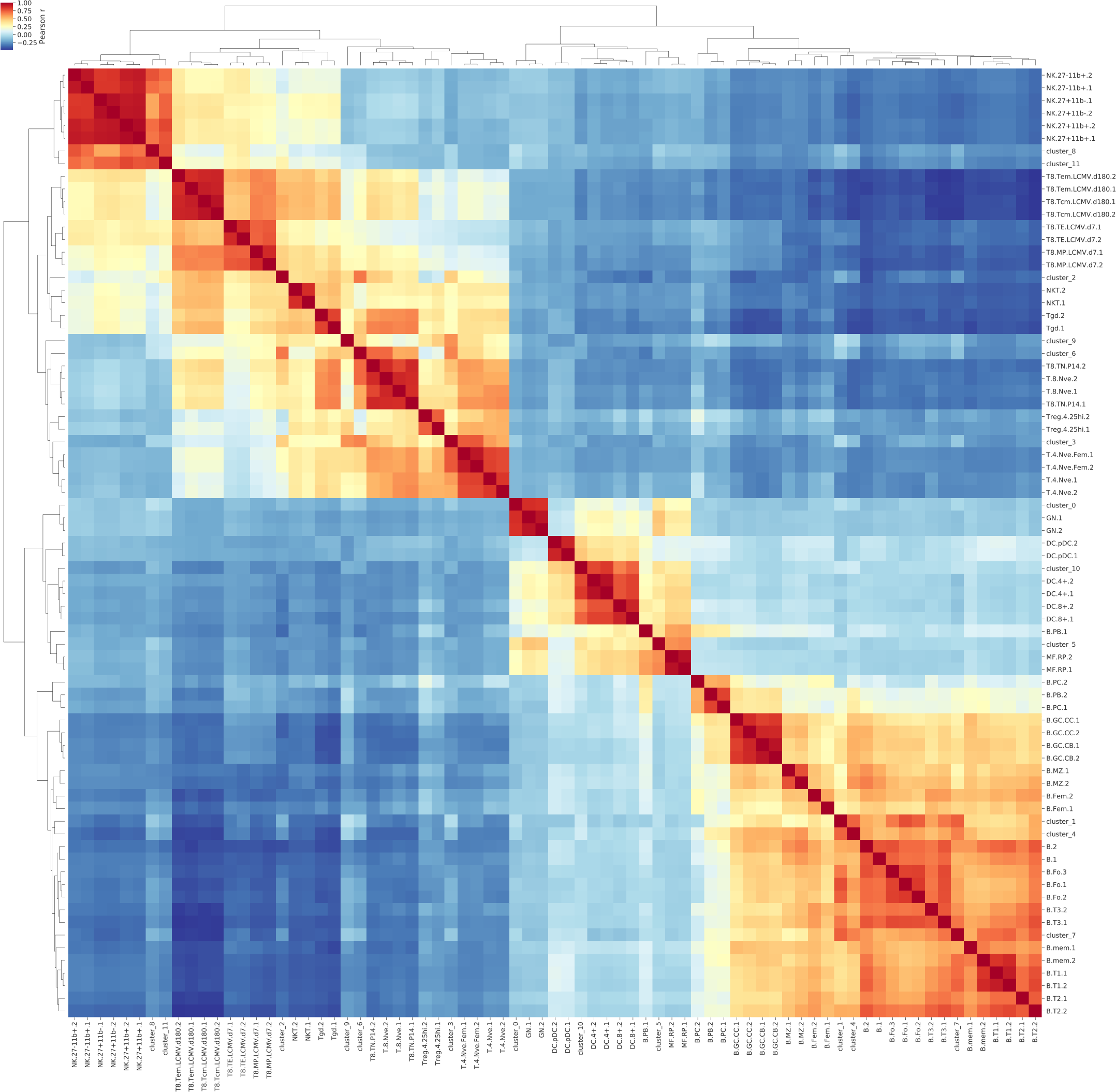
Hierarchical clustering of the Pearson’s correlation between aggregated single cell clusters and the bulk ATAC-seq data sets from ImmGen. The full matrix is shown here, and the ImmGen sample labels were taken directly from the ImmGen ATAC-seq data deposited at the European Nucleotide Archive (ENA) (https://www.ebi.ac.uk/ena/data/view/PRJNA392905).

**Supplementary Figure 7.**
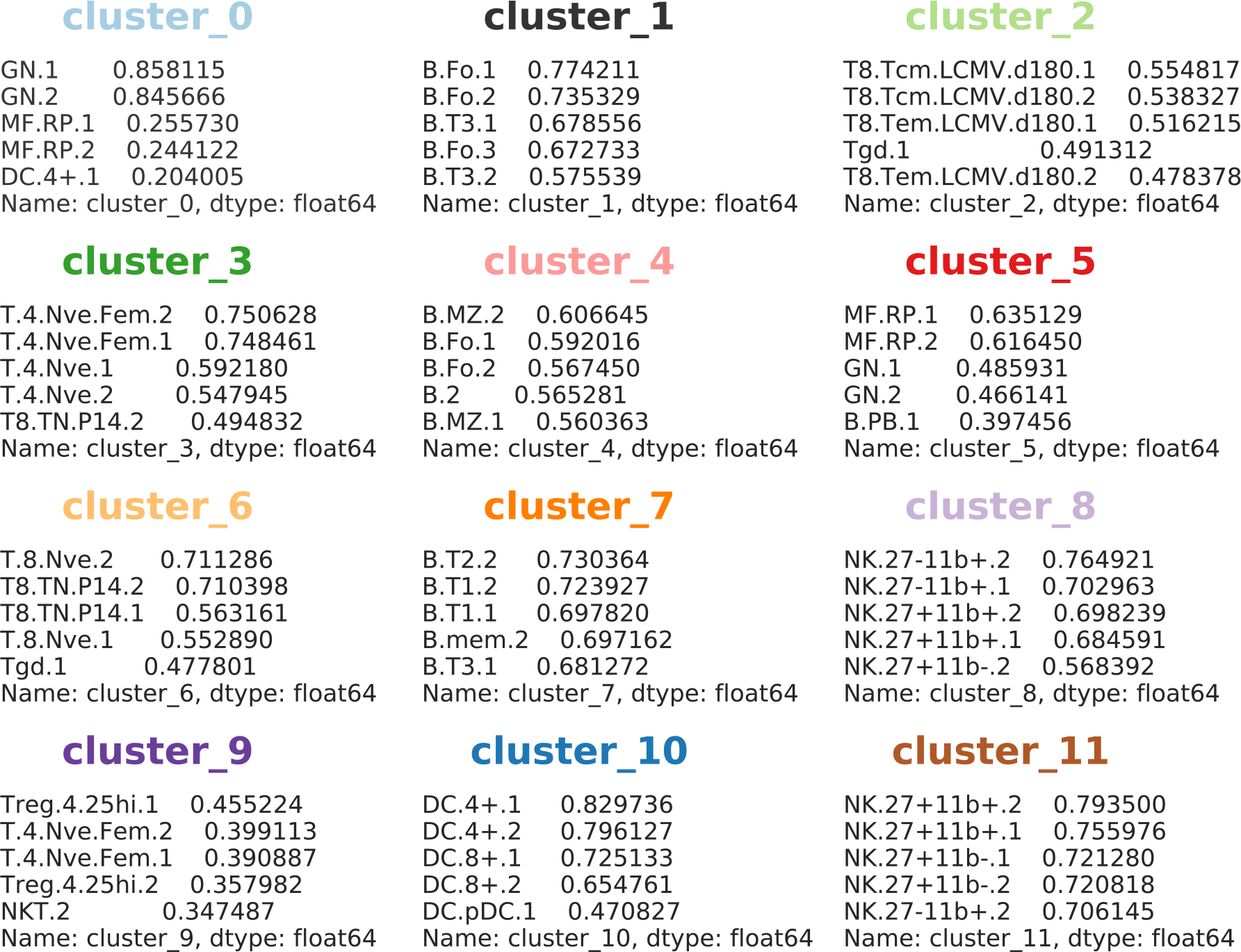
The top correlated ImmGen bulk samples to each aggregated single cell clusters. Top 5 pearson r scores for each clusters are shown.

**Supplementary Figure 8.**
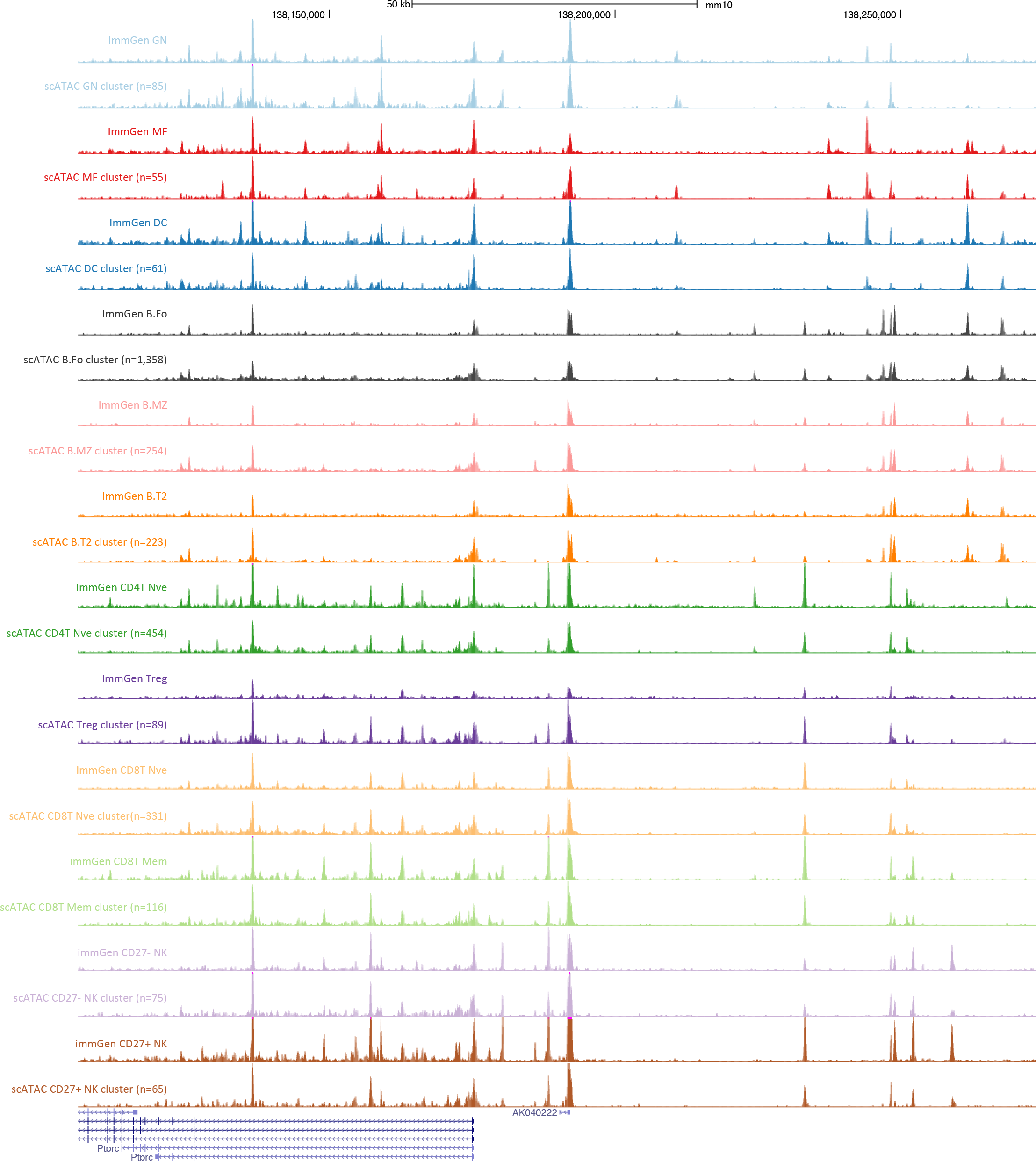
UCSC genome browser tracks showing ATAC-seq profiles of indicated ImmGen bulk samples and aggregated single cell clusters around the *Ptprc* (*Cd45*)promoter region.

**Supplementary Figure 9.**
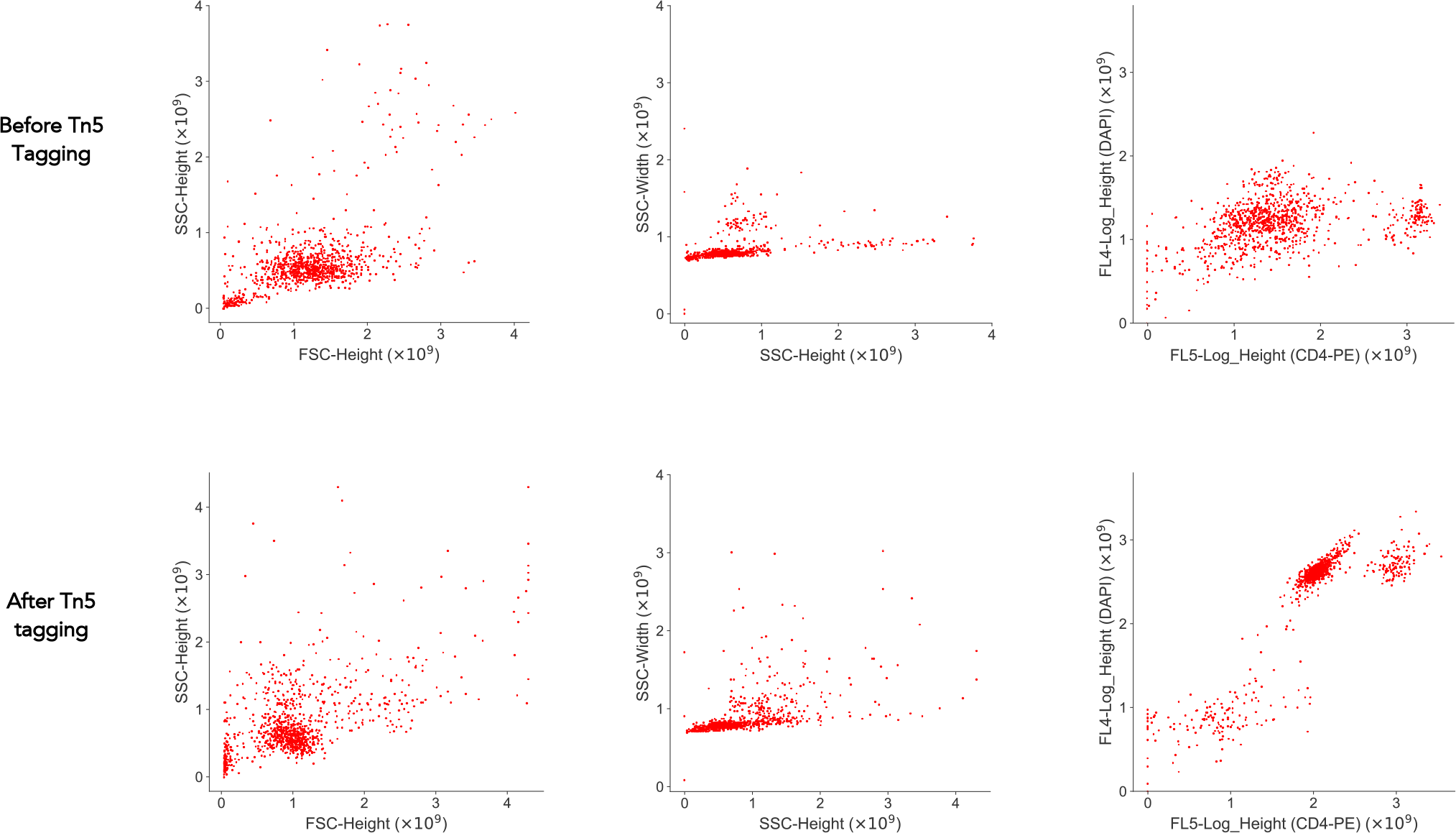
FACS results showing the anti-CD4-PE and DAPI stain on mouse splenocytes before (top) and after (bottom) Tn5 tagging. Note, all cells are DAPI negative before Tn5 tagging but become DAPI positive afterwards. CD4-PE signal remains after Tn5 tagging.

**Supplementary Figure 10.**
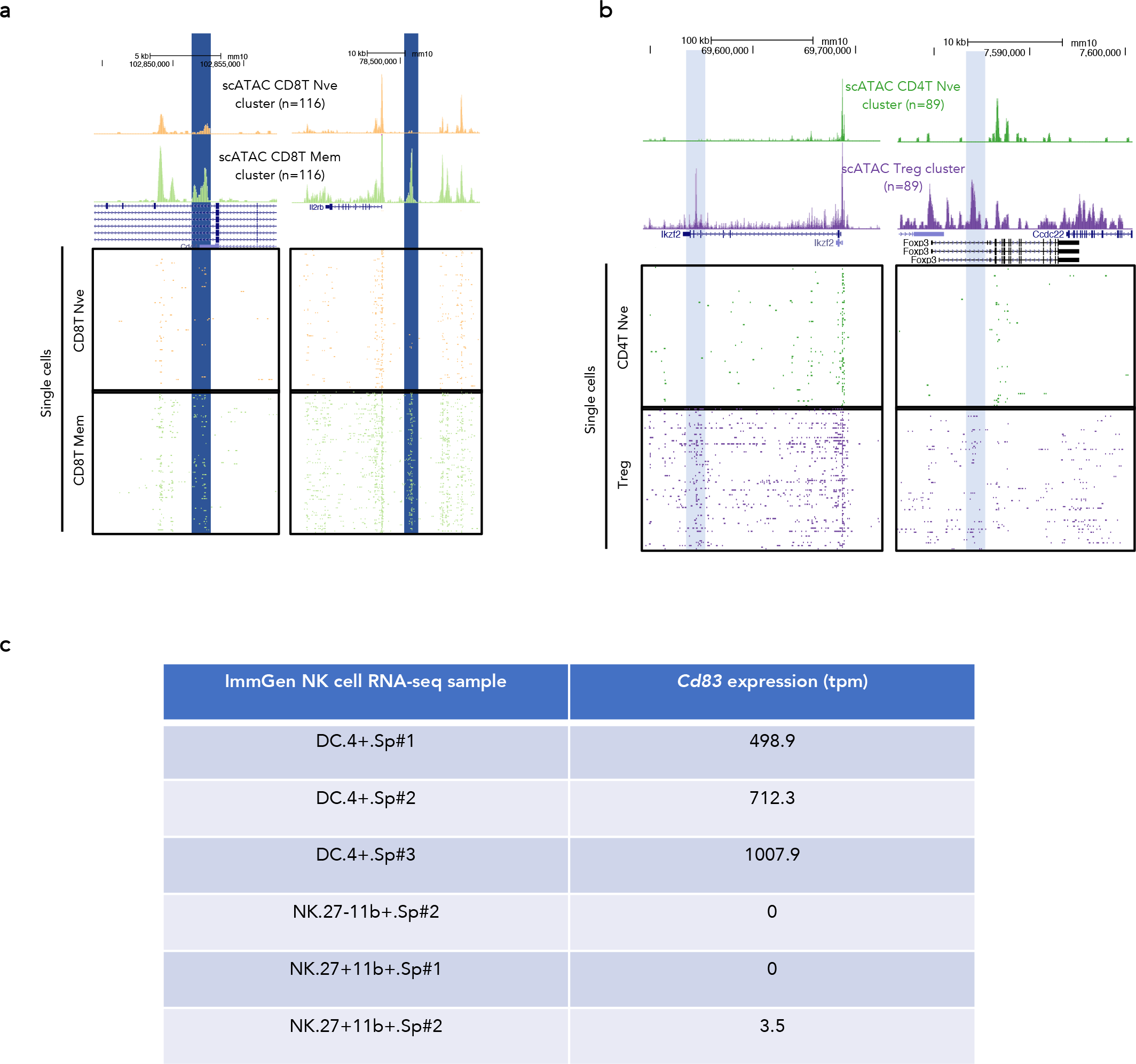
(a and b) UCSC genome browser tracks showing ATAC-seq profiles of aggregate (top panel) and individual single cells (bottom panels). Known enhancers are highlighted. (c) *Cd83* expression from the ImmGen bulk RNA-seq of the indicated sample.

**Supplementary Figure 11.**
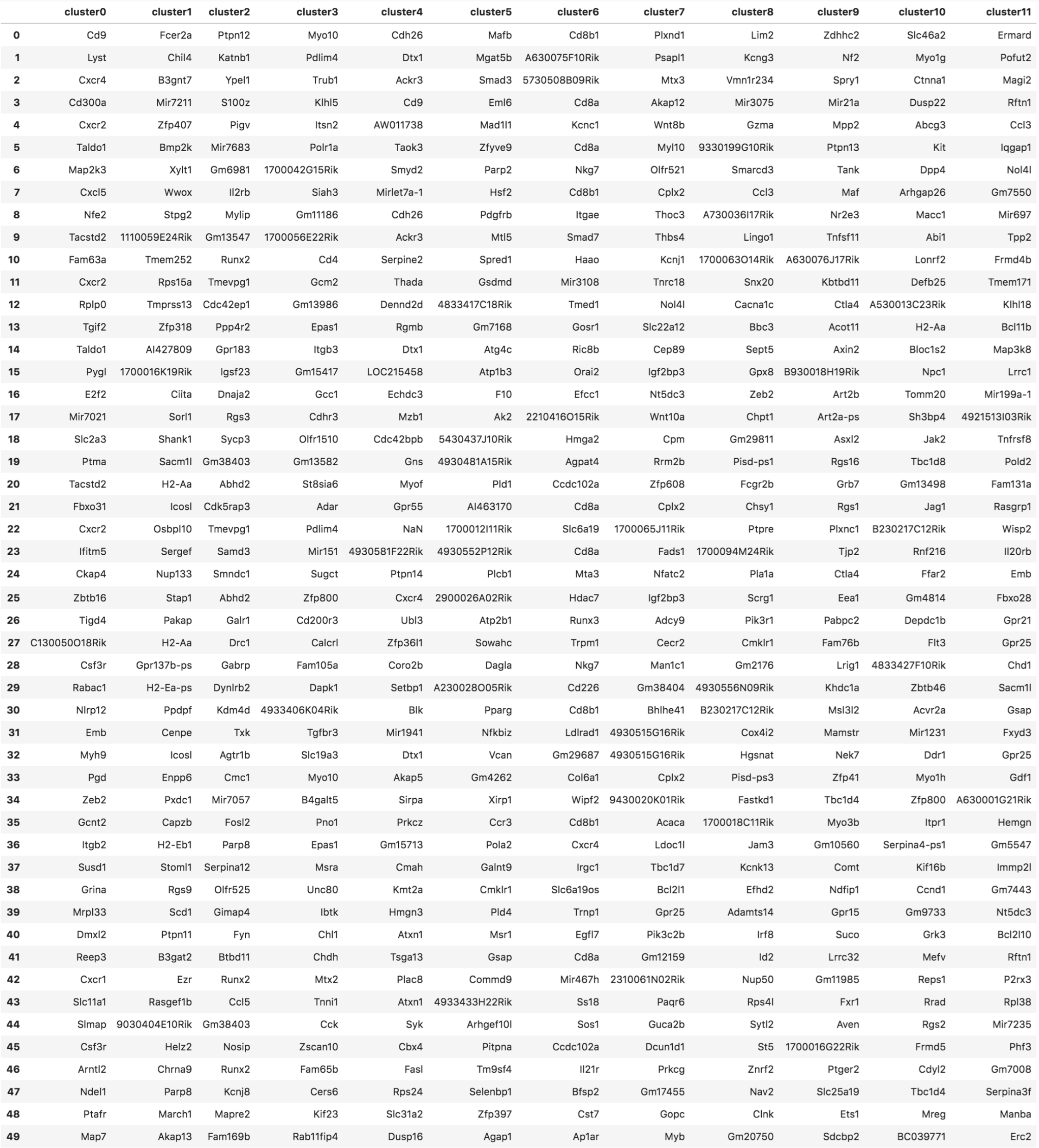
Nearest genes assigned to the top 50 marker peaks in each single cell cluster.

**Supplementary Figure 12.**
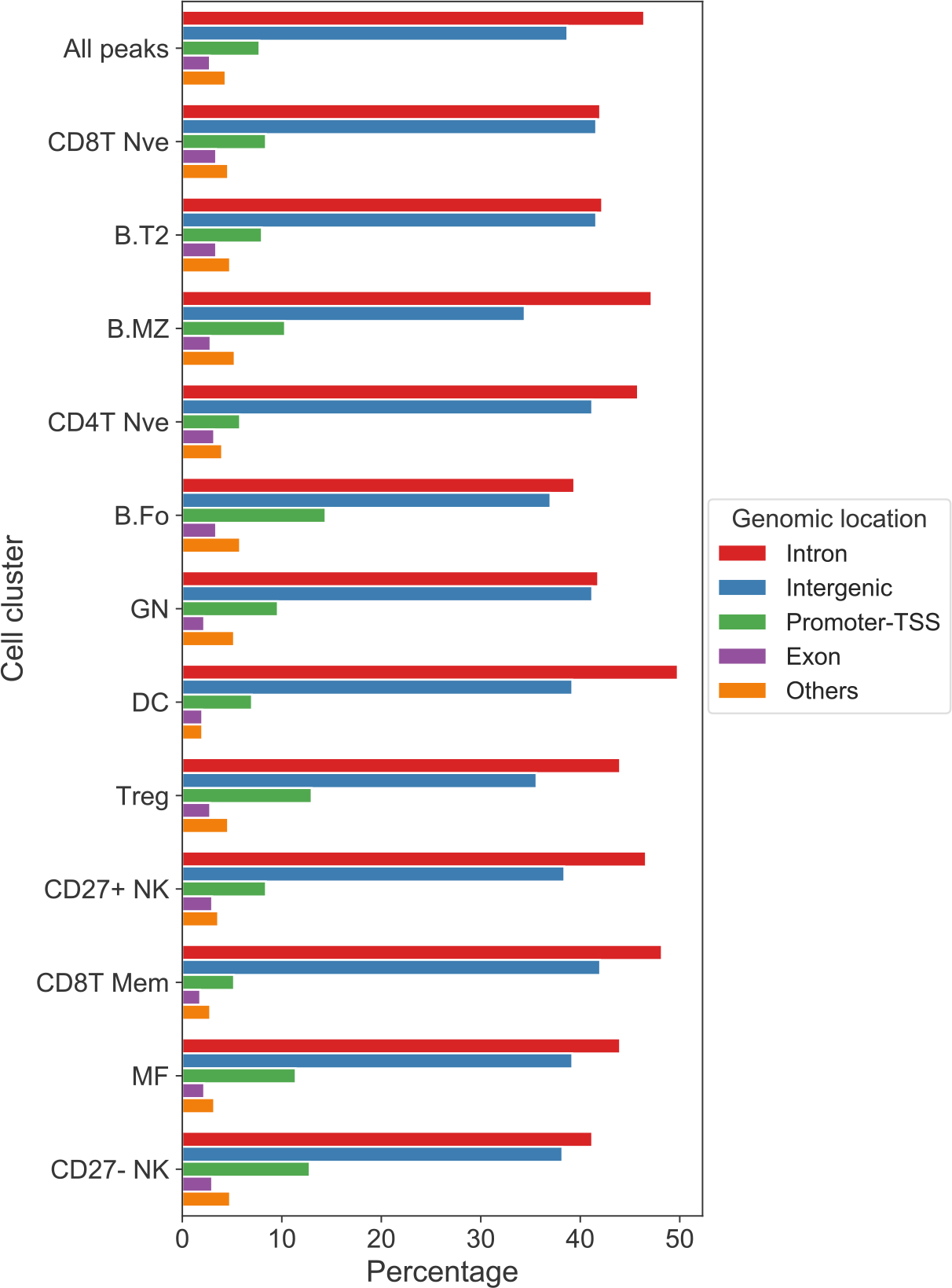
Genomic distribution (by HOMER) of all peaks and the marker peaks in each single cell cluster.

## Supplementary Protocol

Protocol for plate-based scATAC-seq using FACS Timestamp: 15-Feb-2018

1. One day before the experiment, prepare the plates by aliquoting 2 μl 2X Lysis Buffer to each well of the plates (either 96-well or 384-well plate). Then add 2 μl of 10 μM S5xx/N7xx Nextera Index Primer Mix (5 μM each) to each well. Seal the plate and store in −80 °C. Recipe for 2X Lysis Buffer:

2. On the day of the experiment, thaw plates at room temperature.
3. Pre-coat all tubes with 500 μl 0.5% BSA (prepared in 1X PBS) for a few minutes to reduce sample loss. Count or sort 5k – 50k cells into 1.5-ml eppendorf tubes. DO NOT use DNA LoBind tubes for pelleting cells, which does not work well especially when cell numbers are limited.
4. Pellet cells at 500 g, 4 °C, 5 minutes.
5. Wash the cell pellet with 100 μl ice-cold PBS, twice, 500 g, 4 °C, 5 minutes, and carefully remove the supernatant.
6. Resuspend the cell pellet in 50 μl tagmentation mix. The recipe for the tagmentation mix is as follows (THS-seq recipe):

|  |  |
| --- | --- |
| 100 mM | Tris.HCl, pH 8.0 |
| 100 mM | NaCl |
| 40 μg/ml | Proteinase K (Ambion, AM2546, 20 mg/ml stock) |
| 0.4% | SDS |

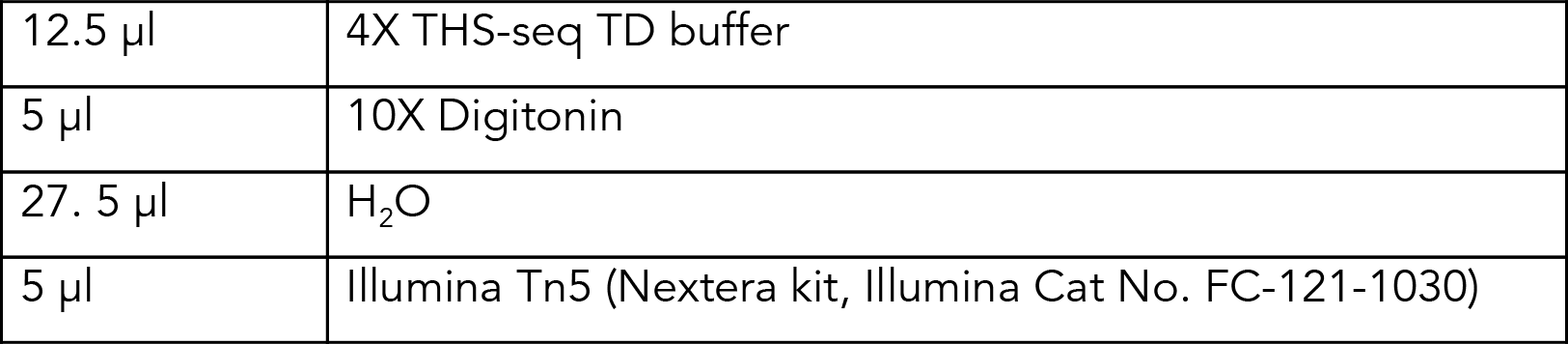 Recipe for 4X THS-seq TD buffer:

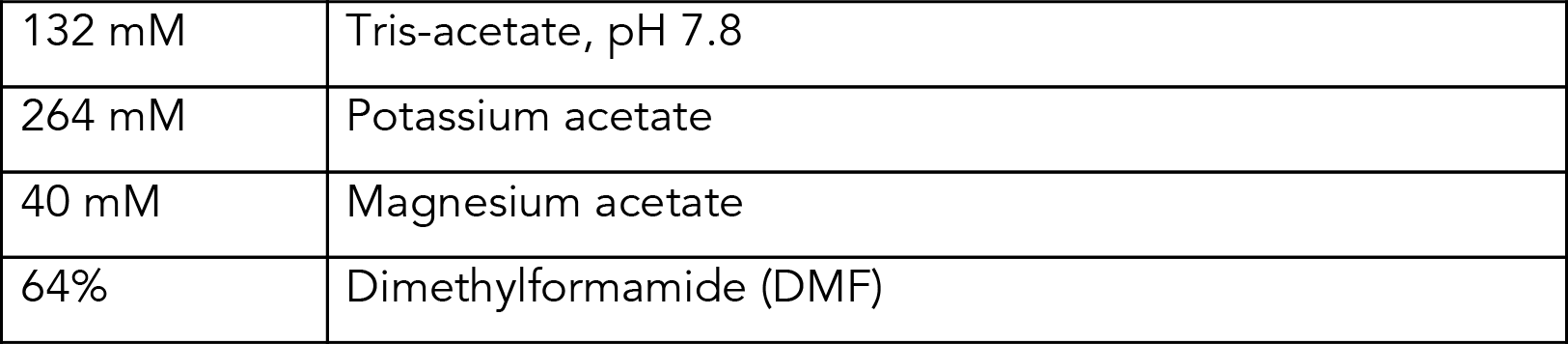 Recipe for 10X Digitonin:

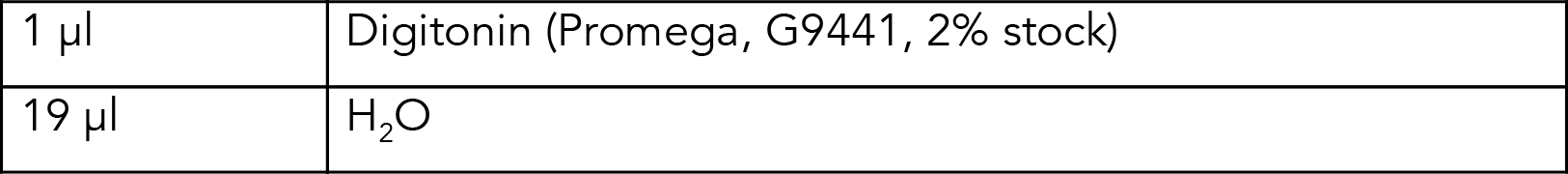
7. Put the tagmentation reaction (50 μl) on a thermomixer, 37 °C, 800 rpm, 30 minutes.
8. Stop the reaction by adding 50 μl tagmentation stop buffer (TSB). Recipe for TSB:

Recipe for 10X Digitonin:
|  |  |
| --- | --- |
| 1 μl | Digitonin (Promega, G9441, 2% stock) |
| 19 μl | H<sub>2</sub>O |

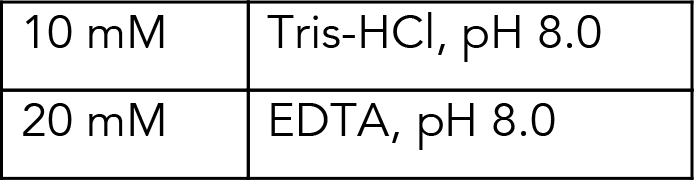
9. Leave on ice for 10 minutes.
10. Add 100 – 300 μl PBS/0.5% BSA to the 100 μl stopped tagmentation mix, and transfer to a FACS tube.
11. Optional: add DAPI to stain nuclei based on manufacturer’s instruction.
12. Sort DAPI positive single nuclei into the plates prepared the day before.
13. Quickly spin down and seal the plate well (can be stored in −80 °C for a few weeks from here), and put the plate on a PCR machine, with lid temperature set to 100 °C.
14. Incubate the plate at 65 °C for 15 minutes to perform Tn5 release.
15. Add equal volume (4 μl) of 10% TWEEN-20 to each well to quench SDS. Briefly vortex to mix.
16. Add 2 μl H_2_O to each well.
17. Add 10 μl 2X NEBNext^®^ High-Fidelity 2X PCR Master Mix (NEB M0541L) to each well
18. At this stage, each well contains 20 μl PCR reaction.
19. Perform library amplification PCR: 72 °C 10 minutes 98 °C 5 minutes [98 °C 10 seconds, 63 °C 30 seconds, 72 °C 20 seconds] × 18 10 °C hold
20. Combine all reactions into a 50-ml falcon, which yields about 20 μl × 384 = 7.68 ml. Normally, the yield will be ~ 7.2 ml.
21. Add 5 volumes (~ 36 ml) Buffer PB (Qiagen), mix well, and pass reaction volume through a single column from a Qiagen MinElute PCR Purification Kit by connecting the column to a vacuum.
22. To wash the column, pass through 40 ml Column Wash Buffer (10 mM Tris-HCl, pH 7.5, 80% ethanol).
23. Spin down the column at top speed on a table top centrifuge to remove all traces of ethanol, and remember to use a pipette to remove the ethanol leftover on the rim of the Qiagen column.
24. Elute the library in 12.5 μl Buffer EB. Perform the elution three times and combine the three elutes to a final volume of ~ 36 μl.
25. Do a final fragment size selection using 0.5X SPRI upper cutoff, followed by 1.2× SPRI lower cutoff, and elute in 30 μl 10 mM Tris-HCl, pH 8.0.
26. Run Nanodrop to obtain a rough estimate of the concentration, and then dilute the library to a range suitable for Bioanalyzer/TapeStation etc.
27. Check for expected results (see Supplementary Fig. 2a).
28. Sequencing: we sequenced each 384 pool on one lane of Hiseq 2000 or one rapid run of Hiseq 2500, which nearly saturated the library. From the data obtained, each cell was sequenced to about 1 million reads, but only ~30,000 unique reads were obtained per cell. Further reads were redundant, which is comparable (if not better) to published scATAC-seq by other methods. Theoretically, 30,000 reads per cell should be sufficient to profile the unique reads. However, considering the presence of mitochondrial DNA, non-mapped and non-uniquely mapped reads, it is safer to aim for at least 100,000 reads per cell.

Oligonucleotides sequence

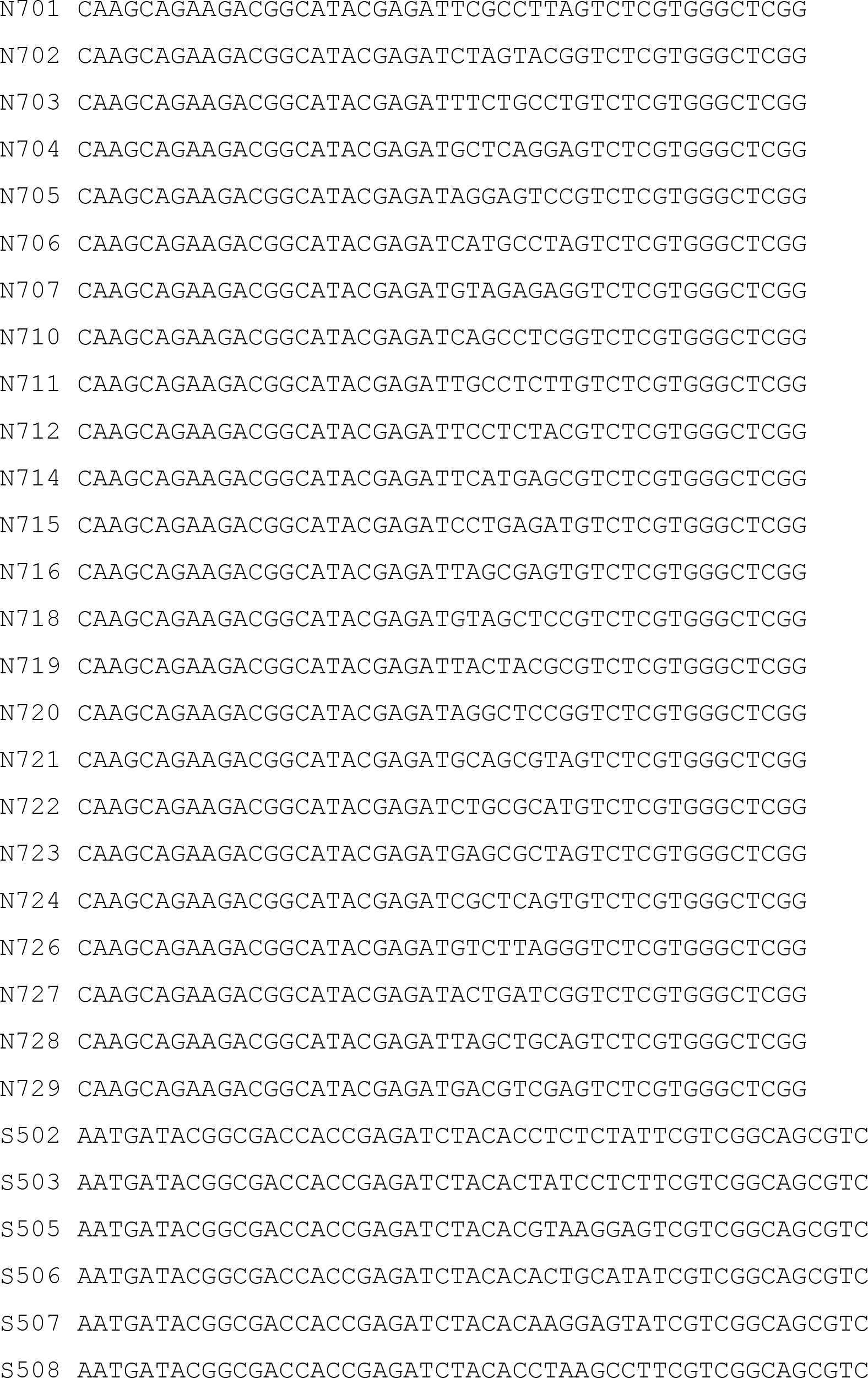

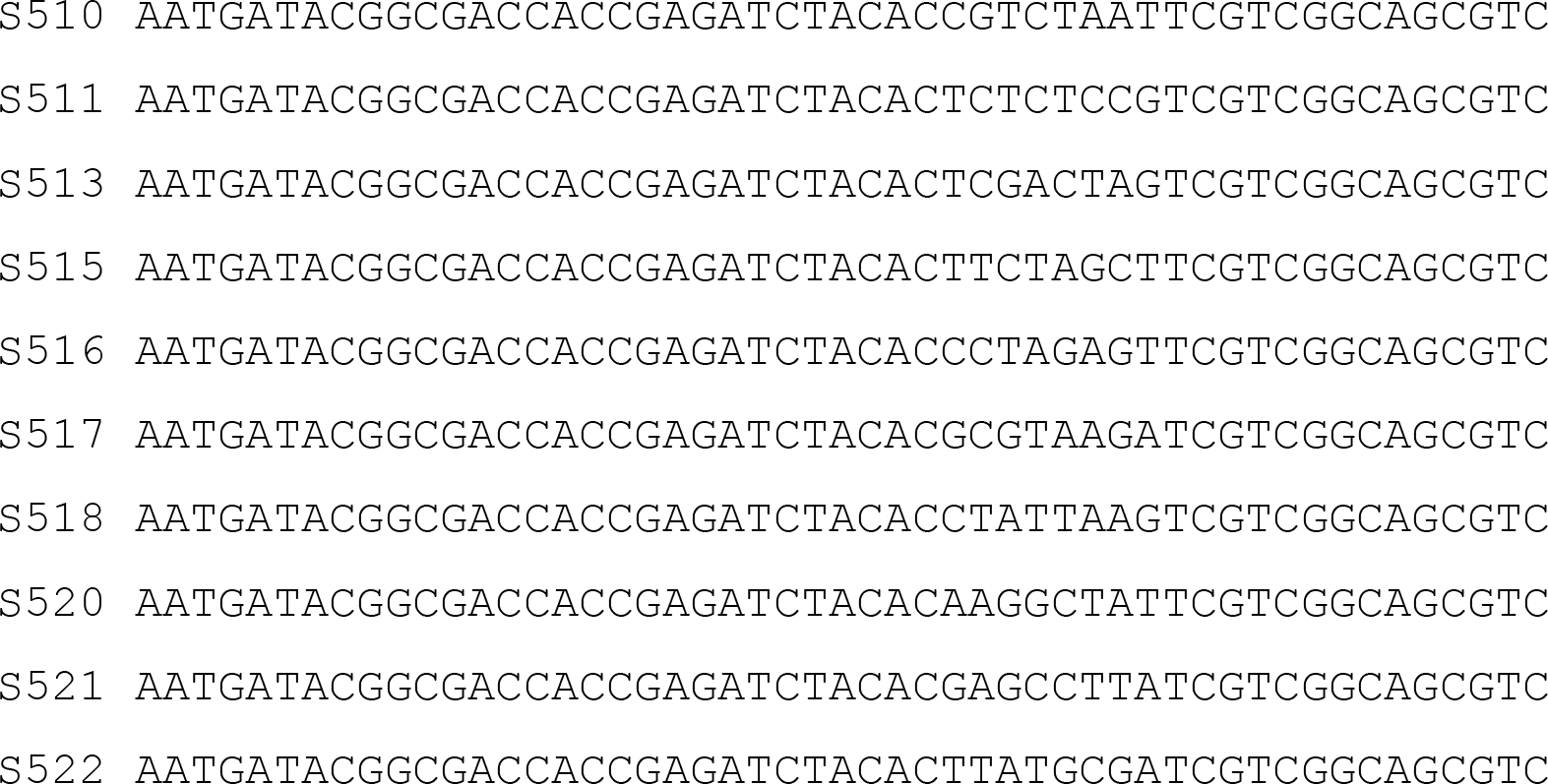

